# Genomic expansion in the dark: A repeat-rich genome of a cave-dwelling remipede sheds light on the evolution of the hexapod sister group

**DOI:** 10.64898/2026.10.30.762620

**Authors:** Elizabeth Borda, Prashant P. Sharma, Carlos E. Santibáñez-López, Jesús A. Ballesteros, Aarón García-Bernal, Fernando Calderón-Gutiérrez, Nuno Simões, Beatriz Yáñez-Rivera, Raúl Llera-Herrera

## Abstract

Remipedes are enigmatic blind, venomous crustaceans endemic to anchialine cave ecosystems (aphotic, hypoxic subterranean environments) and occupy a pivotal phylogenomic position within Pancrustacea as the sister group to Hexapoda. A high-quality genome assembly has long proven elusive for this taxon, due to accessibility challenges and the fragility of remipedes when removed from their habitat. Here we present a highly contiguous, chromosome-level genome assembly for the remipede *Xibalbanus tulumensis* (Xibalbanidae) spanning 8.05 Gb on 30 chromosome-scale scaffolds, placing it among the largest arthropod genomes reported to date. The protein-coding gene repertoire was highly conserved, with over 97% of arthropod orthologs recovered, and microRNA surveys showed a high level of completeness (94%) relative to pancrustacean genomes. Genome expansion appears to be driven primarily by proliferation of intronic and repetitive elements, which comprise ∼71% of the genome. Comparative genomic analyses reveal extensive gene family turnover, with relatively few families showing lineage-specific expansions and contractions. Venom-associated gene families included the remipede-specific Xibalbins, while significant expansion and episodic positive selection affected other venom-associated gene families. Comparative analysis of macrosynteny revealed that the remipede genome retains the signature of ancient fusion-with-mixing events that unite Pancrustacea and Allotriocarida, with considerable similarity in the chromosome architecture of remipedes and basally branching insect orders. Our findings provide the first genomic framework for Remipedia and reveal a highly expanded, repeat-rich genome that retains deep phylogenetic signals in its architecture, despite being adapted to a highly unusual environment.

## Introduction

Extreme environments provide natural laboratories for investigating genome evolution under strong ecological and physiological constraints (Goh et al. 2023; Rappaport and Oliverio 2023). Species inhabiting environments characterized by chronic environmental stress, resource limitation, and ecological isolation exhibit diverse genomic signatures of adaptation, including gene loss, gene family turnover, transposable element proliferation, and lineage-specific functional innovation (McGaugh et al. 2014; Kim et al. 2019; Wang et al. 2019; Yuan et al. 2022; Zhang et al. 2025). Such patterns have been documented in species found across diverse ecosystems, including polar regions (Kim et al. 2019; Lee et al. 2024), hypersaline lakes (Chen et al. 2025), deep-sea ecosystems (Wang et al. 2019; Thomas-Bulle et al. 2022; Xu et al. 2023; Hilali et al. 2025), and subterranean environments (McGaugh et al. 2014; Recknagel and Trontelj 2022; Warren et al. 2024).

Anchialine ecosystems, also known as subterranean estuaries, are karstic flooded cave systems characterized by salinity stratification, variable oxygen availability, complex biogeochemical cycling, chemosynthetic activity, and sulfide-rich groundwaters (Pohlman et al. 1997; Pohlman 2011; Humphreys et al. 2015; Brankovits et al. 2017; van Hengstum et al. 2019; Brankovits and Pohlman 2020). As such, these habitats support specialized faunas that exhibit high levels of endemism and evolutionary novelty (Yager 1981; Iliffe and Kornicker 2009; Neiber et al. 2012; Humphreys et al. 2015; Ballou et al. 2022; Calderón-Gutiérrez et al. 2024). Among the most enigmatic inhabitants of anchialine systems are remipedes, a small clade of blind crustaceans restricted to saline, often hypoxic groundwaters and distributed disjunctly across the Caribbean, Canary Islands and Western Australia (Yager 1981; Koenemann et al. 2007; Neiber et al. 2011; Neiber et al. 2012; Hoenemann et al. 2013; Koenemann and Iliffe 2013; Yager 2013; Olesen et al. 2017; Gonzalez et al. 2020). Remipedes possess a highly distinctive body plan consisting of a cephalic shield followed by an elongated trunk bearing 30−42 pairs of homonomous swimming appendages; they are hermaphrodites that produce lecithotrophic naupliar larvae (Koenemann et al. 2009; Neiber et al. 2011; Koenemann and Iliffe 2013). Unlike many subterranean crustaceans that occupy detrital and grazing niches, remipedes are predators that occupy upper trophic levels within anchialine food webs (Pohlman et al. 1997; Koenemann and Iliffe 2013). They are also unique among crustaceans in possessing a venom delivery system used for prey capture, making them the only known venomous crustaceans to date (von Reumont et al. 2014a; von Reumont et al. 2014b). Remipedes also occupy a pivotal phylogenomic position within Pancrustacea as the sister group to Hexapoda, representing an ancient lineage that diverged from insects more than 450 million years ago (von Reumont et al. 2012; Schwentner et al. 2017). This placement is difficult to reconcile with paleontological data, as no anatomical traits outright support the remipede-hexapod relationship and it also raises questions about the terrestrialization of hexapods. One proposed scenario places the last common ancestor of remipedes and hexapods in shallow marine environments, followed by independent colonization of anchialine habitats by remipedes and terrestrial environments by hexapods (von Reumont et al. 2012). Separately, transcriptomic, morphological, and protein-level studies have provided important insights into remipede venom composition and function (von Reumont et al. 2014a; von Reumont et al. 2014b; von Reumont et al. 2017).

Investigations such as these would be greatly facilitated by high-quality genomic resources for remipedes. However, the absence of genome assemblies for the group has hindered broader investigations of genomic architecture, gene family evolution, and the genomic basis for lineage-specific innovations. This is partly attributable to the challenge of collecting suitable remipede tissue (and especially live specimens) from their habitat for the purposes of long read sequencing and Hi-C scaffolding, as remipedes are highly fragile when exposed to environmental changes. Here, we present the first chromosome-level genome assembly for the remipede *Xibalbanus tulumensis*, a species endemic to the Yucatan Peninsula, Mexico. We applied comparative genomic and macrosyntenic analyses to examine patterns of genome evolution in a highly specialized subterranean pancrustacean. This resource is anticipated to unlock new dimensions of inquiry for understanding the evolution of Pancrustacea and Hexapoda.

## Results and Discussion

### Chromosome-level assembly and selected gene family completeness

We generated a high-quality *de novo* genome assembly for the remipede *Xibalbanus tulumensis* (***Figure 1; Video S1***) using PacBio HiFi long-read sequencing, providing the first chromosome-level genomic resource for the pancrustacean class Remipedia. The primary assembly produced a highly contiguous genome spanning approximately 8.05 Gb at 17x, with a relatively low heterozygosity rate ranging between 0.14−0.45% (***Table S1***). To enable chromosomal scale scaffolding, chromatin contact data were generated using Omni-C library preparation and sequencing using three whole-body samples preserved in ethanol. Incorporation of long-range contact information substantially improved genome contiguity (***Figure 2a***), producing a final scaffolded assembly composed of 3,351 scaffolds ≥500 bp, with a scaffold N50 of 259 Mb, a largest scaffold of 600.4 Mb, and L50 of 12 scaffolds. The assembly had a GC content of 38.1% and contained few ambiguous bases (5.88 Ns per 100 kbp) introduced during scaffolding. Genome completeness was assessed using BUSCO v5.8.2 (Simão et al. 2015; Manni et al. 2021) with the arthropoda_odb10 dataset, recovering 97.9% complete BUSCOs, the majority of which were present as single copy (97%), with low duplication (0.9%) and few fragmented (1.9%) or missing (0.2%) orthologs (***Figure 2b***). Multiple near-complete and partial mitochondrial genome scaffolds (12.9−17.9 kb; 98.7−99.1% similarity) were recovered from the assembly through Blastn searches using the published *Xibalbanus (=Speleonectes) tulumensis* mitochondrial genome as a reference (Lavrov et al. 2004). Searches also identified several mitochondrial-derived fragments on the chromosome-scale scaffolds (i.e., Scaffolds 2, 4, 8, 24), ranging from 3.6−9.1 kb and exhibiting 76− 97% similarity to the mitochondrial genome reference, potentially consistent with candidate nuclear mitochondrial insertions (NUMTs) (***Table S2***).

**Figure 1.**
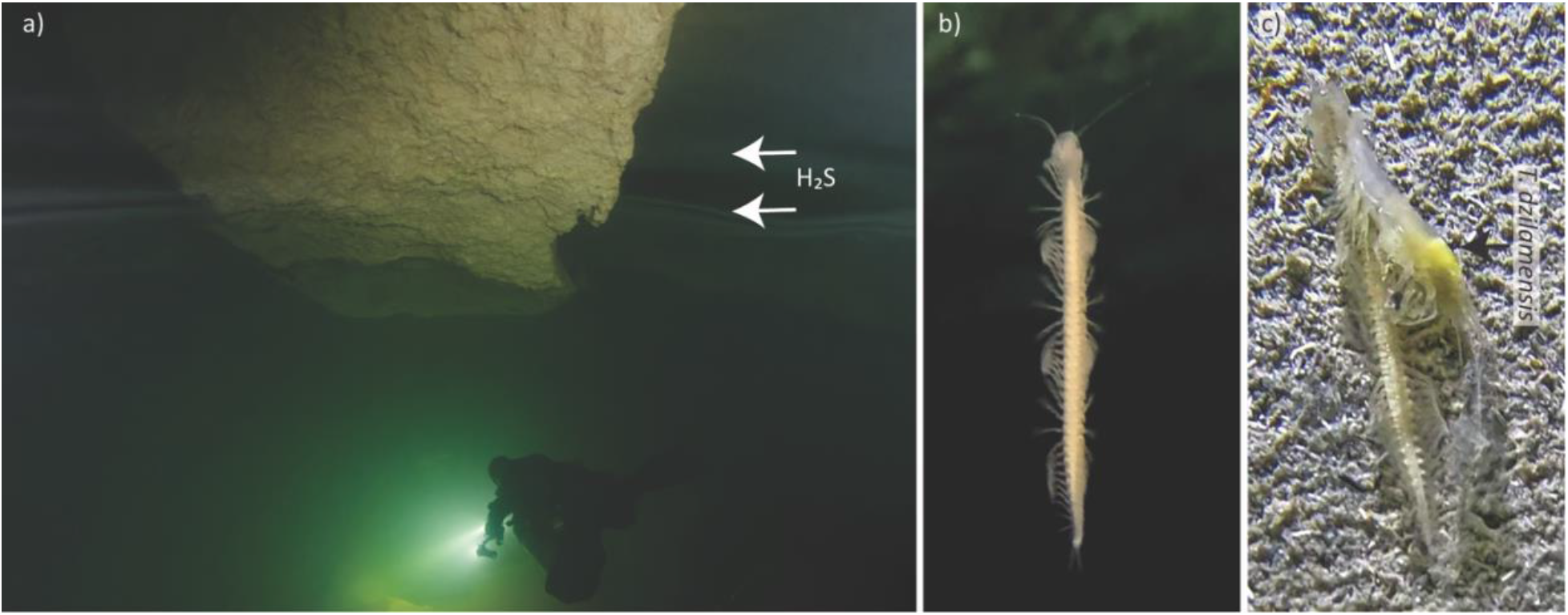
Habitat and behavior of the remipede *Xibalbanus tulumensis*. **a)** Habitat of *X. tulumensis* in the cave section with high density; downstream from cenote Crustacea (Aayin Aak system). Arrows indicate presence of hydrogen sulfide at the halocline. In situ video photograms of *X. tulumensis*: **b)** swimming in the water column; and **c)** with a captured cave shrimp *Typhlatya dzilamensis*. Photo credits: (a) Vlada Dekina; (b−c) Fernando Calderón Gutiérrez. See also **Video S1**.

**Figure 2.**
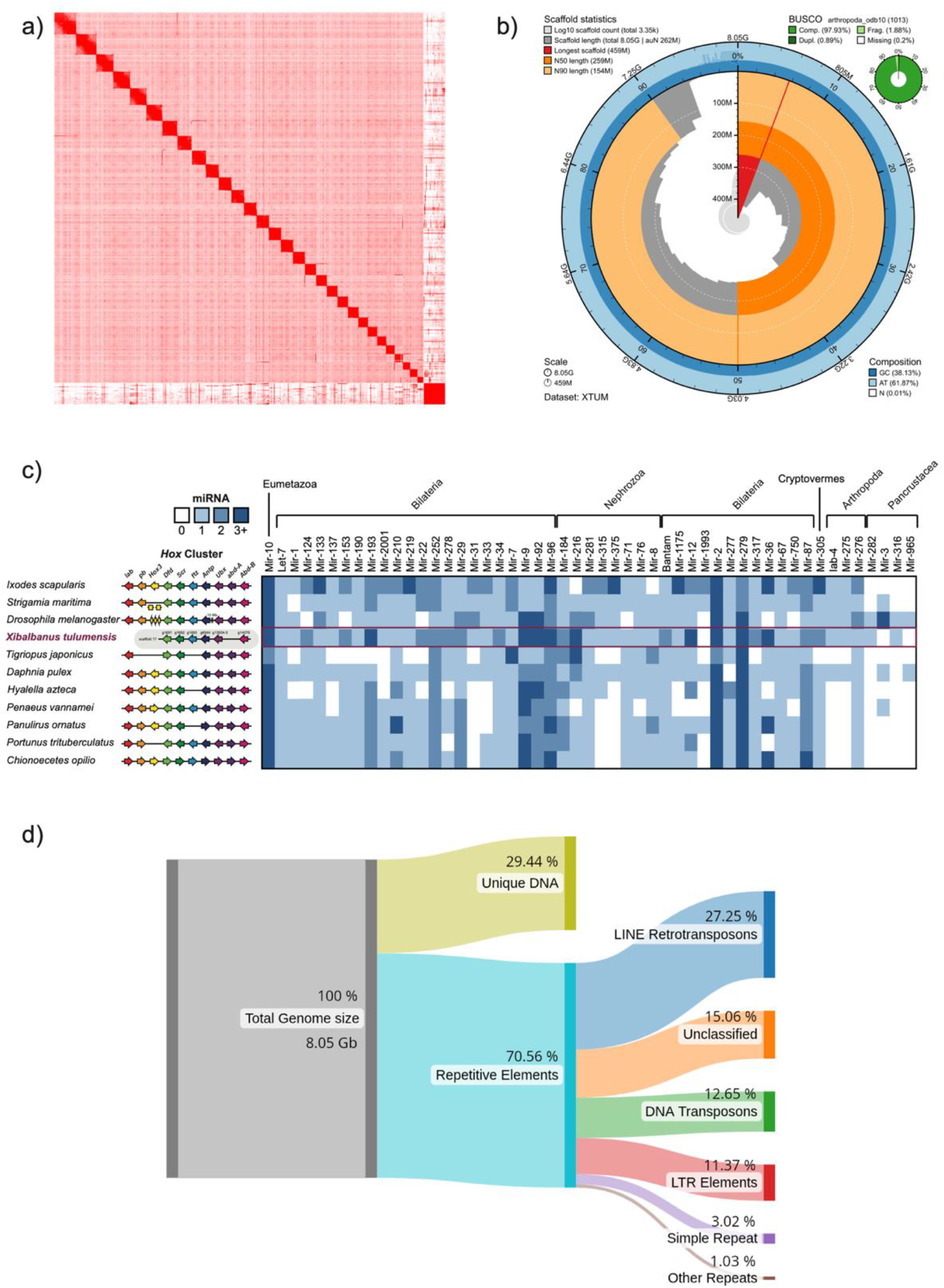
Chromosome-scale genome assembly conserved developmental and miRNA architecture, repetitive landscape and functional gene distribution in *Xibalbanus tulumensis*. **a)** Hi-C contact map of the final assembly showing ∼30 chromosome-scale scaffolds. **b)** Snail summarizing assembly statistics and BUSCO completeness (C:97.9% [S:97.0%, D:0.9%], F: 1.9%, M: 0.2%, n=1013). The genome spans 8.05 Gb, with a contig N50 of 258 Mb and the largest scaffold of ∼600 Mb. **c)** (left panel) Comparative organization of Hox gene clusters across select pancrustacean taxa. Colored arrows indicate the relative order and transcriptional orientation of Hox genes. While most taxa retain a conserved, colinear Hox cluster, Xibalbanus tulumensis exhibited a fragmented arrangement across scaffold 17, with several Hox genes not recovered in the current genome assembly. (right panel) Heatmap of miRNA copy numbers per family (0, 1, 2, 3+ copies) showing conserved metazoan miRNA is broadly retained in *X. tulumensis*. **d)** Sankey diagram showing repetitive element composition revealing repeat-driven architecture (70.6%), dominated by LINE retrotransposons, followed by unclassified repeats, DNA transposons, LTR elements and minor fractions of simple and other repeats.

To assess completeness with respect to conserved gene families and clusters, we examined selected gene repertoires comparatively across pancrustacean taxa (***Table S3***), revealing strong conservation of core regulatory architectures. Analysis of developmental gene families revealed a conserved arrangement of *Hox* genes (***Figure 2c***, left panel), despite the absence of some cluster members (e.g., *lab, pb, Hox3,* and *abd-A*). The involvement of several of these *Hox* genes in essential arthropod head-patterning functions, as well as their consistent presence in other arthropod genomes, suggests incomplete sequencing of the cluster, rather than true *Hox* gene losses (Gainett et al. 2023; Nicolini et al. 2023). Next, we explored the repertoire of microRNA (miRNA) families (***Figure 2c***, right panel) using the mirMachine pipeline (Umu et al. 2023), as miRNA complements are reliably conserved as single-copy except in cases of whole-genome duplication (Leite et al. 2016; Ontano et al. 2021; Gainett et al. 2024). Out of 51 miRNAs expected in typical pancrustacean genomes, homologs of 48 were identified in the *X. tulumensis* genome and these did not exhibit evidence of systemic duplication compared to other crustacean exemplars (***Figure 2c***, right panel; ***Table S4***). Notably, the miRNA families considered to be putatively Pancrustacea-specific (*mir-3*, *mir-282*, *mir-316*, *mir-965*) were the least well represented across the 11 arthropod genomes surveyed. *mir-3* was identified in the genome of the centipede *S. maritima*, whereas *mir-282* and *mir-316* were found only in the genomes of *X. tulumensis* and the hexapod exemplar *D. melanogaster*. These results suggest that the genome of the remipede is fairly complete and does not exhibit the signature of whole-genome duplication.

### Genome architecture reveals extreme repeat-driven expansion

Repetitive elements constitute a substantial fraction of the *X. tulumensis* genome, accounting for 70.6% of the assembled sequence and representing the primary contributors to its large genome size (***Figure 2d***). RepeatMasker analyses showed a high abundance of transposable elements (TEs) (***Table S5***), with long interspersed nuclear element (LINE) retrotransposons comprising the dominant class (27.3% of the genome), including major contributions from the RTE-BovB, CR1, and Penelope families (∼1.9 Gb combined). Long terminal repeat (LTR) retrotransposons also represented a significant component of the genome (11.4%), with Gypsy and unclassified LTR elements together accounting for ∼907.5 Mb. DNA transposons contributed an additional 12.7% of genomic content. In addition to TEs, simple repeats and low-complexity regions together account for ∼3.5% of the genome, and a substantial fraction of repetitive sequences were unclassified (15.1%), consistent with lineage-specific or highly diverged repeats. These results support that the expanded genome size of *X. tulumensis* is driven primarily by repeat proliferation, as reported in other pancrustacean taxa (Alfsnes et al. 2017; Hultgren et al. 2018), rather than increases in gene number, large-scale duplication or polyploidy.

### Protein coding gene content and genome organization

Protein-coding gene annotations identified 16,698 gene models comprising 24,110 transcripts in the *X. tulumensis* genome (***Table S6***). Despite the exceptionally large genome size (∼8.05 Gb), gene number and exon organization fall well within the range reported for other arthropods (Thomas et al. 2020), supporting conservation of core gene content. Coding regions are compact (mean CDS length: 1 kb), whereas gene lengths are substantially expanded (mean: 86.0 kb), consistent with extensive intronic expansion. In *X. tulumensis*, genome size inflation is decoupled from increases in gene number, reflecting expansion of non-coding genomic components.

Among predicted protein sequences, 1,418 (∼5.9%; ***Figure S2***) lacked significant BLAST hits but were assigned conserved domain and GO annotations by InterProScan (Jones et al. 2014), suggesting functional conservation despite substantial sequence divergence. An additional 5,443 transcripts (∼22.5%) contained identifiable protein domains but lacked GO annotations, indicating limited functional resolution or poorly characterized domain families. These proteins were distributed across numerous low-frequency domain families, reflecting broad functional diversity among highly divergent protein sequences.

Functional annotation based on GO terms and conserved domains identified canonical gene repertoires associated with ion transport, oxidative stress responses, environmental sensing and signaling, reproduction and toxin-/venom-related functions (***Figure S1*; *Table S7***). These categories were examined because of their expected relevance to the physicochemical conditions of anchialine environments, including salinity gradients and hypoxia (Gorr et al. 2006; Humphreys et al. 2015; Brankovits et al. 2017; Brankovits and Pohlman 2020), and to aspects of remipede biology, including reproduction, sensory biology, and venom function (Yager 1981; Koenemann et al. 2009; Koenemann and Iliffe 2013; von Reumont et al. 2014a). While some functional categories were diverse, including genes associated with osmoregulation (n=205), genes associated with oxidative stress and detoxification (n=62) and reproduction (n=12) were less represented. Additionally, despite neuroanatomical evidence that remipedes possess highly structured sensory and olfactory centers (Koenemann and Iliffe 2013; Stemme et al. 2016), only 88 genes associated with environmental sensing and signaling were recovered, and extensive diversification of this repertoire was not observed. In contrast, genes associated with toxin- and venom-related enzymatic functions (n=521) were highly diverse. Transcript-level annotations were distributed across the chromosome-level scaffolds, with individual scaffolds harboring between 7 and 38 annotations. Proteolytic and enzymatic components such as serine endopeptidases and metallopeptidases exhibited localized genomic clustering across multiple scaffolds, consistent with patterns observed for the curated venom-specific loci (see ***Figures 5– 6***).

Separately, examining a case of expected gene loss, we searched the genome for homologs of opsins (***Figure 3***), as it had previously been reported that the transcriptome of this species lacks opsins (Henze and Oakley 2015), whereas two opsins (a neuropsin and a long wavelength-sensitive opsin) had been reported in the transcriptome of the Caribbean remipede *Lasionectes entrichoma* (Palecanda et al. 2022). We additionally assembled and surveyed the transcriptomes of three other remipede species from the Bahamas (*Pleomothra apletocheles, Morlockia williamsi*, and *Godzilliognomus frondosus*), for opsins as part of this analysis. We found one pteropsin present in *X. tulumensis* and no additional opsins in the transcriptomes of the remaining three species. These results suggest that degradation and losses of opsins in the anophthalmic Remipedia may have occurred in lineage-specific patterns.

**Figure 3.**
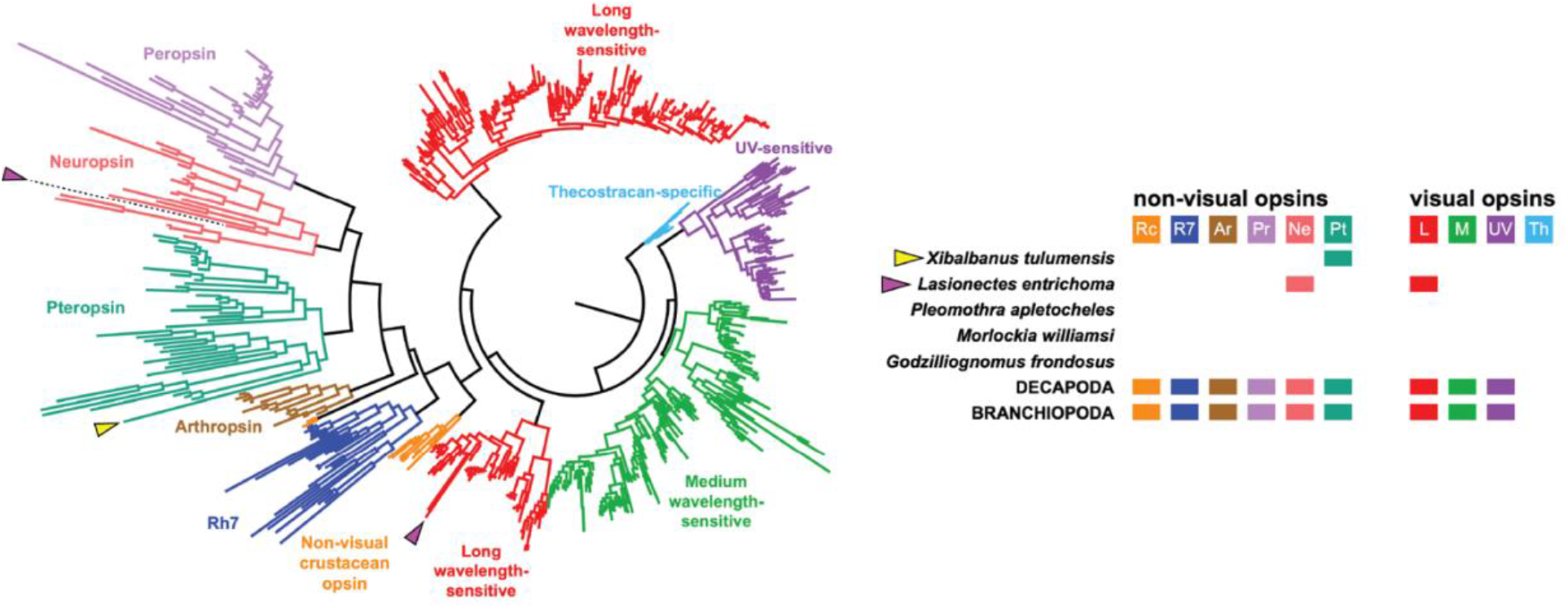
Phylogenetic distribution of non-visual and visual opsin genes across Remipedia, by comparison to Decapoda and Branchiopoda ground plans. Recovered opsin-related sequences in X. tulumensis were restricted to one pteropsin homolog in *X. tulumensis* (non-visual); the remipede *Lasionectes entrichoma* was previously reported to possess one neuropsin (non-visual) and long wavelength-sensitive opsin. No opsins were recovered in the transcriptomes of the remipedes *Pleomothra apletocheles*, *Morlockia williamsi*, and *Godzilliognomus frondosus*.

### Functional annotation reveals conserved pathway architecture

KEGG annotations recovered 310 pathways spanning major functional categories (***Table S8***; ***Figure S3***). However, functional gene repertoires appeared to be broadly conserved across categories (***Figure S3***). For example, core components of the hypoxia-inducible factor (HIF) signaling pathway and associated metabolic processes were recovered (***Figure S5–S6***), including key regulators of oxygen sensing (e.g., EGLN/PHD and VHL), as well as downstream genes involved in glycolysis and anaerobic metabolism (e.g., hexokinase, phosphofructokinase, and lactate dehydrogenase), suggesting a conserved cellular response to hypoxia (Gorr et al. 2006). Overall, pathways involved in hypoxia tolerance, stress responses, and environmental sensing are well represented and showed no obvious departures from their canonical composition.

### Evolutionary Dynamics of Gene Families

We characterized gene family evolution across 22 pancrustacean taxa using OrthoFinder, identifying 29,289 Hierarchical Orthologous Groups (HOGs; ***Figure 4a***). The number of lineage-specific HOGs inferred in the *X. tulumensis* genome (N=461) was comparable to other pancrustacean counterparts. Following stringent filtering, 19,662 HOGs were retained for downstream analysis using CAFE 5 (Mendes et al. 2021) and mapped onto a fossil-calibrated phylogeny (Wolfe and Hegna 2014; Cornetti et al. 2019) (***Figure 4b***). Seven distinct models were evaluated, among which the Poisson root distribution with two gamma-rate categories (K=2) was favored (***Figure S7***). Across models, we inferred 652 gene family expansions and 2,607 contractions along the *X. tulumensis* lineage relative to the most recent common ancestor of Remipedia and Hexapoda (***Figure 4b***). However, after applying statistical thresholds, only 42 families were significantly expanded and one was significantly contracted (Viterbi p < 0.05), suggesting that most inferred changes reflect background turnover rather than sustained directional shifts. Most evaluated models converged on this core set of significant families (***Figure 4d***) showing consistency across model configurations. Thus, gene family evolution in *X. tulumensis* is characterized by limited large-scale innovation despite substantial genome expansion.

**Figure 4.**
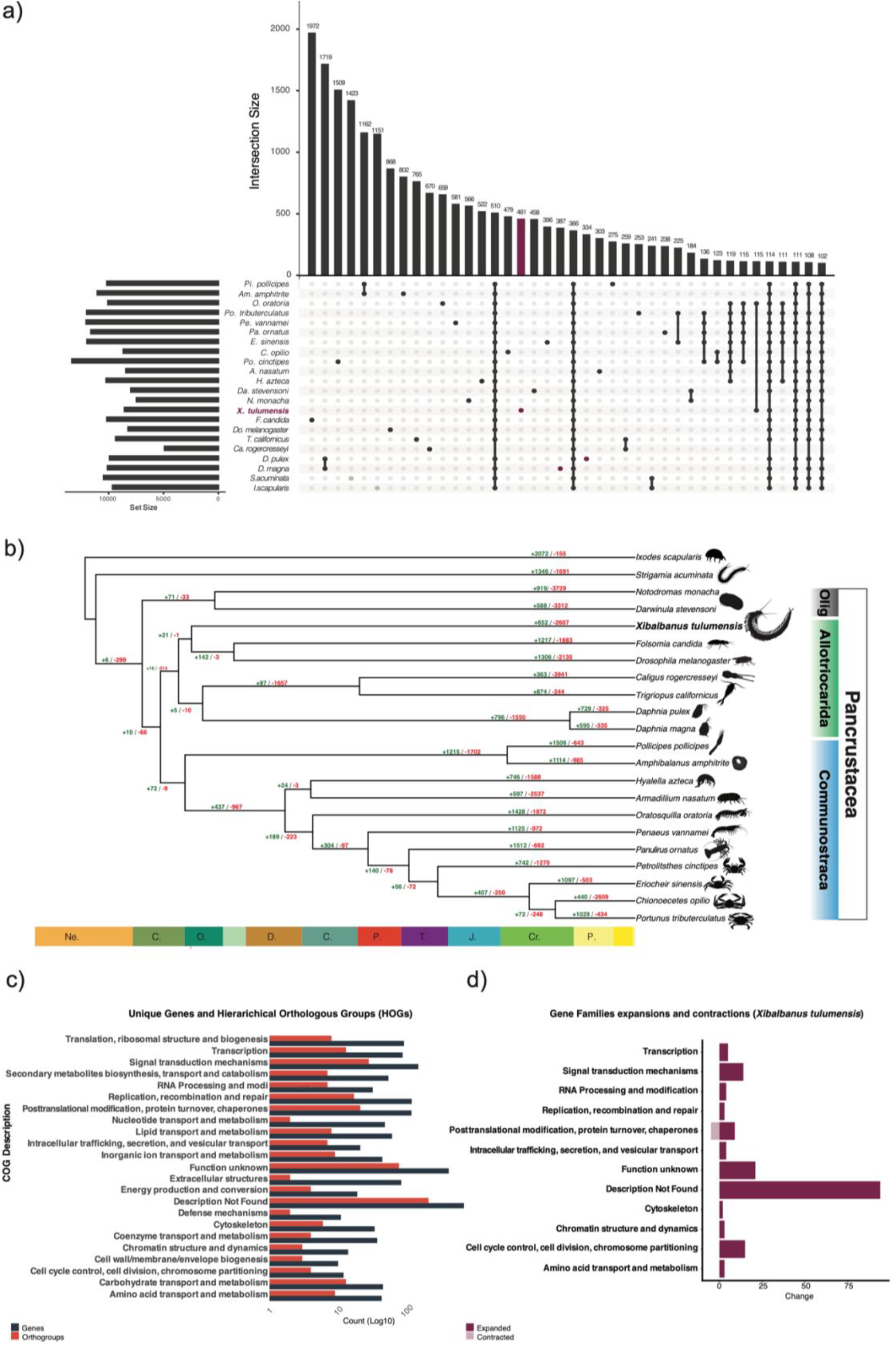
Comparative genomics and gene family dynamics in *Xibalbanus tulumensis* and select Pancrustacea. **a)** Distribution of hierarchical orthologous groups (HOGs) across select pancrustacean taxa inferred from OrthoFinder, showing orthogroup counts and presence-absence patterns across taxa, with *X. tulumensis* highlighted. **b)** Time-calibrated phylogeny with gene family expansion (green) and contractions (red) inferred using CAFE; numbers at nodes indicate significantly expanded or contracted gene families per branch. **c)** Enrichment patterns of functional categorization of genes (COG; black bars) with orthogroups (red bars). **d)** Functional distribution of significantly expanded (maroon bars) and contracted (pink bars) gene families. Silhouettes in **b)** are from phylopic.org, with the following adapted from photographs using ChatGPT: *Petrolisthes* from an image of *P. violaceus* by J. Antonio Baeza 2016, <u>Wikimedia Commons</u>; used to represent the genus; *Eriocheir sinensis* by Lipatov 1966, <u>Wikimedia Commons</u>; *Chionoecetes opilio* by J. Deeds, U.S. Food and Drug Administration, <u>Wikimedia Commons</u>; *Portunus trituberculatus* by Almandine <u>Wikimedia Commons</u>.

### Evolutionary Dynamics of Remipede Venoms

To characterize the evolutionary history of venom components, putative toxin families previously identified by von Reumont et al. (2017) were localized within the *X. tulumensis* genome assembly and assigned to their respective HOGs (***Figure 5a***). A total of 210 putative venom genes were identified, distributed across 64 HOGs. Among these, five HOGs were lineage-specific (“orphans”) to *X. tulumensis* and were excluded from CAFE analysis, while the remaining 59 HOGs were evaluated for gene family expansions and contractions. CAFE5 (Mendes et al. 2021) analyses identified three HOGs undergoing significant expansion (Viterbi p < 0.05; ***Figure 5a****, red bars*).

**Figure 5.**
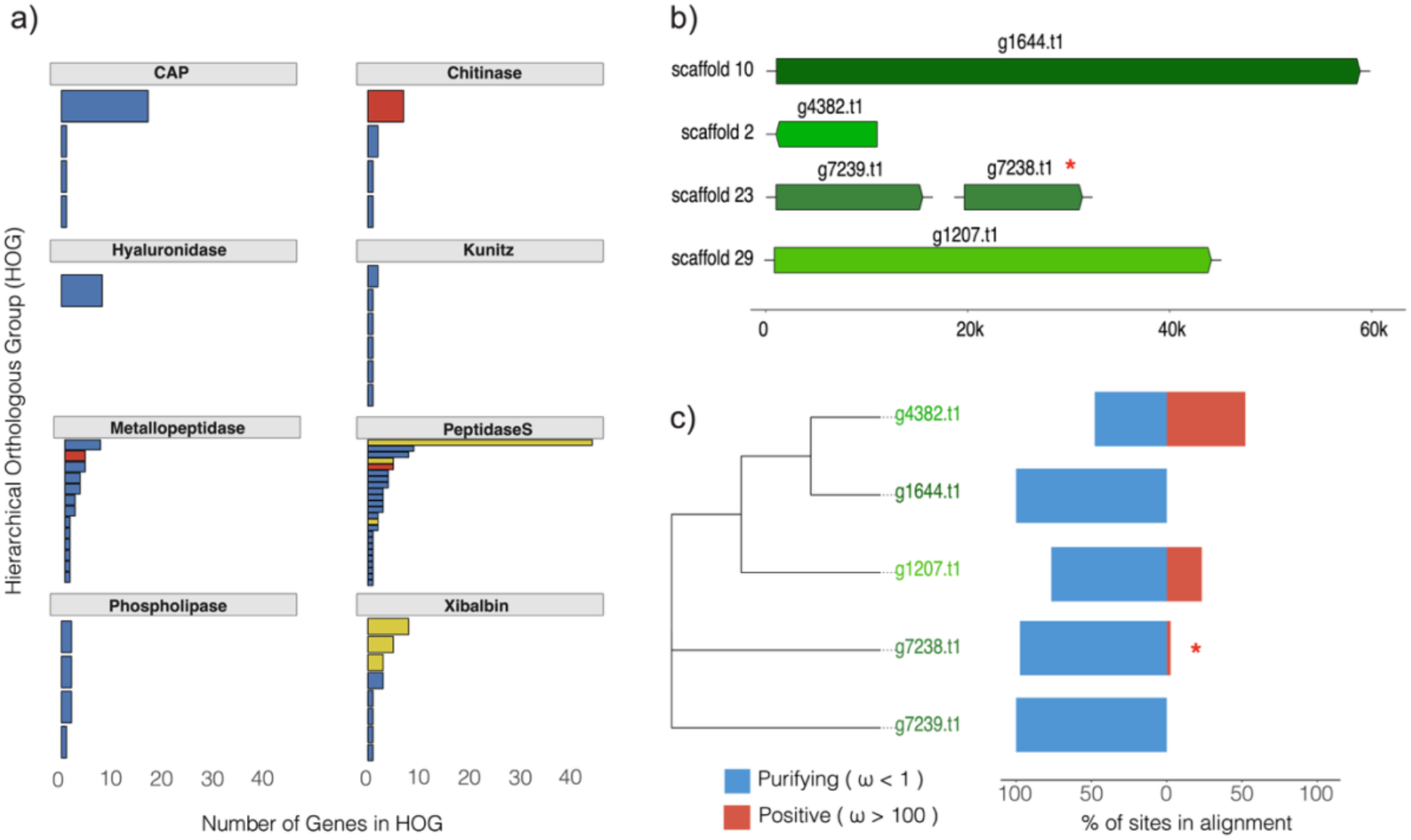
Venom gene family expansion and evolutionary dynamics in *Xibalbanus tulumensis.* **a)** Gene counts across Hierarchical Orthogroups (HOGs) for major venom-associated families, highlighting the expansion of serine peptidases and chitinase proteins. **b)** Genomic distribution of Peptidase S genes from a single expanded HOG (identified via CAFE) across four distinct scaffolds. **c)** Phylogenetic and selection analysis of Peptidase S genes within this expanded HOG, showing most copies under purifying selection (blue, ω < 1), while a subset exhibited strong positive selection (red, ω > 100). Tip label colors correspond to the respective scaffolds shown in **(b)**. expanded by CAFE, which consists of five genes distributed across scaffolds 2, 10, 23, and 29 (Figure 5b).

Putative venom-associated gene families were distributed across eight major functional groups, where gene family sizes varied substantially across toxin classes, including a) Cysteine-rich secretory proteins, Antigen 5, and Pathogenesis-related 1 (CAP), b) Chitinase, c) Hyaluronidase, d) Kunitz, e) Metallopeptidase, f) Peptidase S, g) Phospholipase, and h) Xibalbin (***Figure 5a***). The CAP superfamily proteins comprised 20 genes across four HOGs. Chitinases included 11 genes across four HOGs, with one significantly expanded HOG (n=7 genes). Hyaluronidases were represented by eight genes within a single HOG, while Kunitz-type inhibitors comprised seven genes across six HOGs, including one lineage-specific HOG. Metallopeptidases were represented by 31 genes across 13 HOGs (one HOG expanded significantly containing four genes). Peptidase S family members (110 genes across 24 HOGs) represented the largest toxin classes, with one significantly expanded HOG (with five genes), and three HOGs with 59 genes were identified as lineage-specific. In contrast, phospholipase A2 (PLA2) with seven genes across four HOGs, showed no significant expansion. Lastly, Xibalbins, including 23 genes across eight HOGs, were dispersed across eleven scaffolds (see also ***Figure 5a***). Venom gene families exhibited both tandem clustering and dispersion across the genome. This dynamic is exemplified by a Peptidase S Hierarchical Orthogroup (HOG), identified via OrthoFinder and found to be significantly

To determine whether these gene family expansions were accompanied by adaptive protein evolution, aBSREL analyses were conducted on three significantly expanded HOGs of Peptidase S, Chitinase, and Metallopeptidase, respectively, to detect episodic positive selection. Within the Peptidase S family (***Figure 5c***), aBSREL revealed significant episodic selection in one expanded HOG (g7238_t1; p= 0), where a single sequence showed 3% of sites under strong positive selection (ω>500). Expanded chitinase HOG (***Figure 6a***), one paralog (g8795.t1) exhibited significant signatures of positive selection (p = 0), with 14.4% of sites evolving under strong positive selection (ω > 5388), while the remaining sites (85.5%) were under purifying selection (ω = 0.9607). Chitinase genes were distributed in a tandem repeat in one scaffold (***Figure 6b***).

**Figure 6.**
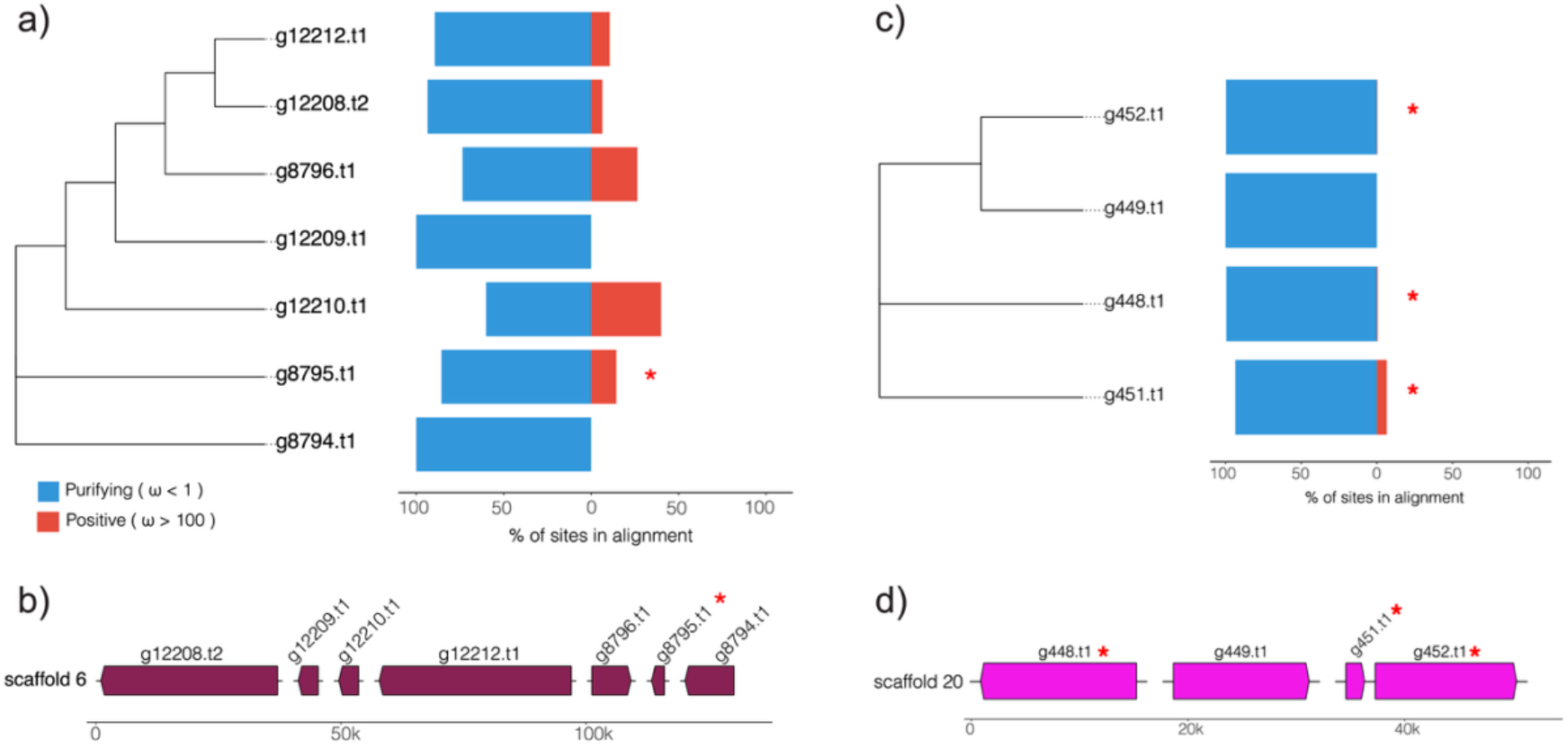
Evolutionary dynamics and genomic organization of additional expanded venom gene families in *Xibalbanus tulumensis*. (a) Phylogenetic relationships of chitinase paralogs from an expanded HOG (identified via CAFE), accompanied by branch-specific selection estimates inferred using aBSREL. (b) Genomic organization of these chitinase genes on a single scaffold (Scaffold 6), exhibiting patterns consistent with tandem duplication. (c) Phylogenetic relationships of metallopeptidase paralogs from an expanded HOG, alongside their aBSREL branch-specific selection estimates. (d) Genomic organization of the metallopeptidase genes on Scaffold 20. Scale bars indicate genomic distance in kilobases (kb). For the phylogenies in (a) and (c), bars indicate the proportion of sites evolving under purifying selection (blue, ω < 1) versus episodic positive selection (red, ω > 100). Asterisks (*) denote branches with significant evidence of positive selection (chitinase: p < 0.0001; metallopeptidase: p < 0.05).

Lastly, within the expanding Metallopeptidase HOG, aBSREL detected significant episodic selection in three of the four transcripts (***Figure 6c***), with 0.3% to 6.5% of sites experiencing positive selection (ω = 33.28, 133.4, and 9583; p = 0.024, 0.031, and 0.020, respectively). These four sequences were in a single scaffold (***Figure 6d***).

### Genome Expansion with Constrained Architecture and Targeted Innovation

Large genome sizes are well documented across arthropods, including the orthopteran *Bryodemella tuberculata* (∼22 Gb) (Hawlitschek et al. 2023), the giant isopod *Bathynomus jamesi* (∼6 Gb) (Yuan et al. 2022) and several caridean shrimp (∼3 − 21 Gb) (Moraes et al. 2025). Genomic expansion in these lineages is driven by the proliferation of repetitive DNA and transposable elements rather than increases in gene number. In comparison with other pancrustacean genomes, including the crayfish *Cherax quadricarinatus* (5.26 Gb) (Chen et al. 2023), the hermit crab *Coenobita brevimanus* (4.74 Gb) (Wang et al. 2024) and the giant freshwater prawn *Macrobrachium rosenbergii* (3.18 Gb) (Zheng et al. 2024), the 8.05 Gb of *X. tulumensis* represents one of the largest arthropod genomes assembled to date. Despite the challenges associated with low biomass and sampling in subterranean environments, the high contiguity of this assembly demonstrates that chromosome-scale genomic resources can be generated for remipedes, with this study providing the first chromosome-scale genomic framework for Remipedia.

Consistent with other arthropods, the exceptionally large genome of *X. tulumensis* is driven primarily by extensive repeat expansion rather than increases in gene content. Despite this genomic inflation, core gene architecture remains broadly conserved, indicating that genome size expansion has occurred without corresponding increases in functional complexity. At the same time, substantial portions of the genome remain poorly characterized, including a large fraction of unclassified repetitive elements, highly divergent protein homologs relative to current databases, and lineage-specific orthologous groups of unknown function. These findings highlight both the evolutionary distinctiveness of Remipedia and the continued underrepresentation of deeply divergent pancrustacean lineages in existing genomic reference resources.

On the other hand, gene family turnover and episodic positive selection were concentrated within a limited subset of functional categories, most notably venom-associated genes (von Reumont et al. 2014a; von Reumont et al. 2014b; von Reumont et al. 2017), highlighting targeted evolutionary innovation within an otherwise conserved genomic framework. Given the central role of venom in prey capture and processing, the expansion and diversification of venom gene families may reflect selection associated with the specialized predatory ecology of remipedes (Pohlman et al. 1997; Carpenter 1999; Koenemann and Iliffe 2013). Unlike the physiological and sensory adaptations often emphasized in studies of organisms inhabiting extreme environments, *X. tulumensis* shows its strongest evidence of lineage-specific innovation within gene families linked to trophic interactions (Pohlman et al. 1997; Koenemann and Iliffe 2013).

### Evolution of Genome Architecture in Pancrustacea

The assembly of a chromosome-scale genome for an exemplar of Remipedia unlocks an understanding of how genome architecture has evolved across Pancrustacea over most of the span of the Phanerozoic. To approach this question, we deployed odp v. 0.3.0 to identify the locations of homologs associated with 29 ancestral linkage groups (ALGs) previously circumscribed for Myriazoa (the Bilaterian-Cnidarian-and-Sponge [BCnS] ALGs (Simakov et al. 2022)). Previous investigations of invertebrate genomes using this suite of tools have shown that various taxa retain these linkage groups with remarkable fidelity (Simakov et al. 2022; Schultz et al. 2023; Lewin et al. 2025). A phylogenetically useful property of these ALGs occurs where chromosomes undergo ancient fusion events, followed by extensive mixing of genes associated with each ALG brought together by the fusion of two chromosomes. Over time, the pattern of mixing is thought to become irreversible, providing a strong phylogenetic character system for testing relationships inferred by other phylogenetic data classes; this approach has been used to reconstruct the placement of Ctenophora in the animal tree of life, as well as to substantiate the monophyly of Lophophorata (Schultz et al. 2023; Lewin et al. 2025).

Various arthropod genomes retain the BCnS ALGs with high fidelity, as previously shown for some members of Chelicerata (Klementz et al. 2025; Kulkarni et al. 2025). However, genome compaction, reduced karyotypes, and accelerated evolutionary rates tend to erase the signature of the ALGs, as exemplified by many acariform mite genomes (Klementz et al. 2025). Such analyses have not been applied across crustacean genomes. We compared the arrangement of BCnS homologs in *X. tulumensis* to those of six other crustacean taxa spanning major splits in pancrustacean phylogeny, as well as to basally branching hexapod exemplars. Despite the extreme environments inhabited by Remipedia and the unusually large genome of *X. tulumensis*, we observed considerable cohesion of the BCnS linkage groups, with 2036 homologs identified across the 29 BCnS ALGs, and 27 out of 29 ALGs exhibiting significant spatial clustering of genes (> 5 genes; p < 0.05) on 29 of the 30 scaffolds of *X. tulumensis* (***Figure 7***; ***Figure S8***). Promisingly, we found similar cohesion of the BCnS ALGs in both crustacean and hexapod exemplars, with two exceptions. The malacostracan *Penaeus vannamei* exhibited no cohesion of the ALGs whatsoever (***Figure S9a***), whereas the branchiopod *Daphnia pulex* exhibited a similar condition, though five ALGs were detected with significance (***Figures S9b***).

**Figure 7.**
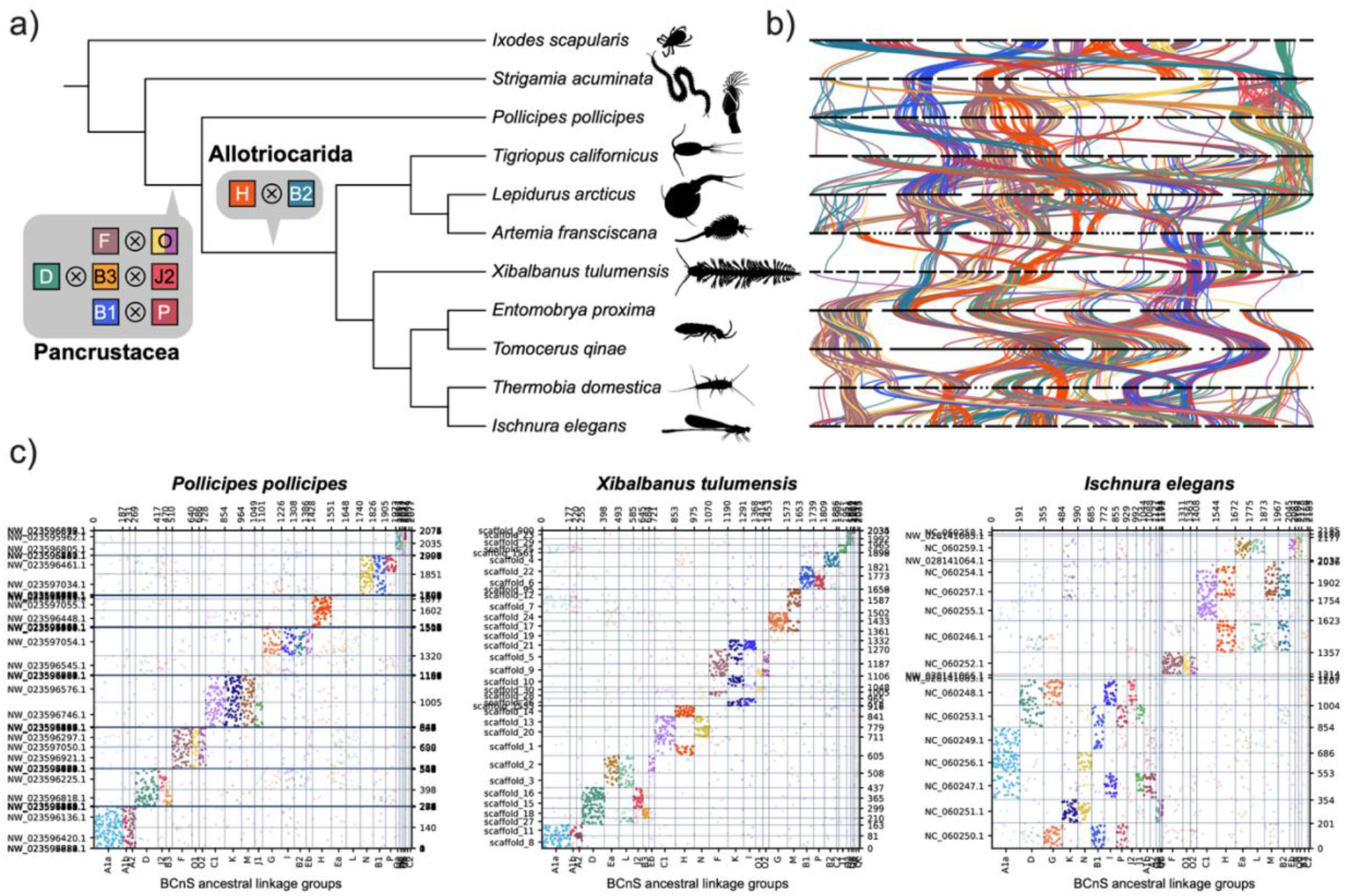
Macrosynteny across the phylogeny of Pancrustacea. **a)** Location of fusion-with-mixing events on a cladogram of Arthropoda. Phylogenetic relationships are based on C60 + LG + F + G maximum likelihood tree of Bernot et al. (2023). Branch lengths are not to scale. Silhouettes are from phylopic.org. **b)** Ribbon plot showing members of the nine ALGs that exhibit fusion-with-mixing. **c)** Oxford dot plots showing the locations of homologs of BCnS loci for the barnacle *Pollicipes pollicipes* (bottom left), the remipede *Xibalbanus tulumensis* (bottom center), and the odonate *Ischnura elegans* (bottom right). Insignificant hits (p > 0.05) have been removed for clarity.

Next, we mapped the incidence of shared fusion-with-mixing events across the phylogeny of Pancrustacea (***Figure 7a***). We identified three fusion-with-mixing events shared by all Pancrustacea, to the exclusion of the myriapod and chelicerate outgroups: F⊗O1⊗O2; D⊗B3⊗J2; and B1⊗P. Due to the absence of chromosome-level genome assemblies for representatives of Oligostraca (the earliest branching clade of Pancrustacea, which includes ostracods, mystacocarids, and Branchiura; Bernot et al. 2023), we cannot rule out the possibility that one or more of these three fusion-with-mixing events are restricted to the non-Oligostraca pancrustaceans. Nevertheless, we observed the signature of these three events in the cirripede exemplar *Pollicipes pollicipes* (Cirripedia and Malacostraca form Communostraca, the second major branch of crustaceans), as well as exemplars of major lineages of Allotriocarida (a clade comprised of copepods, branchiopods, cephalocarids, remipedes, and hexapods), with the caveat that a Cephalocarida genome is also not presently available to reinforce this ancestral state reconstruction.

We additionally observed a fourth fusion-with-mixing event restricted to Allotriocarida (H⊗B2), which informs the placement of Copepoda in the pancrustacean tree of life. Copepoda have historically been unstable across phylogenomic analyses, with site heterogeneous model (CAT-GTR)-based Bayesian analyses alternatively placing them as sister group to Malacostraca within Multicrustacea (Schwentner et al. 2017), as the sister group of Remipedia + Hexapoda within Allotriocarida (Lozano-Fernandez et al. 2019), or as the sister group of Thecostraca within Multicrustacea under Dayhoff-6 state recoding (Lozano-Fernandez et al. 2019). An analytically intensive approach with improved taxon sampling was recently executed (Bernot et al. 2023) which showed that a maximum likelihood approach implementing the per mean site frequency (PMSF) model C60 + LG + F + G recovered Copepoda as the sister group to Branchiopoda within Allotriocarida, whereas CAT-GTR-based Bayesian analyses of the same matrix (with or without Dayhoff recoding) placed copepods as the sister group to Remipedia. These results were shown to be sensitive to taxon sampling, as a remipede-hexapod relationship was only reliably recovered when balancing taxon sampling to include more major lineages of Pancrustacea.

The discovery of a rare genomic change uniting Allotriocarida (i.e., the fusion-with-mixing event H⊗B2, shared by copepods, branchiopods, remipedes, and hexapods) substantiates the conclusions of Bernot et al. (2023) and refutes earlier suggestions that copepods are nested within Communostraca. This consilience between analyses of macrosynteny and the sampling-intensive phylogenomic study of Bernot et al. (2023) also disfavors the reductive interpretation that the CAT-GTR model and/or Dayhoff-6 state recoding represent a panacea for phylogenomic inference (Lozano-Fernandez et al. 2019). New chromosome-scale assemblies of understudied groups such as Cephalocarida, Ostracoda, and Branchiura are anticipated to test these hypotheses further, as well as shed more light on the evolution of pancrustacean genomes.

## Conclusion

In the 14 years that have passed since Remipedia were recognized as the likely sister group of the hexapods, understanding of remipede biology has advanced mainly through transcriptomic datasets. The genome assembly provided in this study, the first of its kind for any remipede, breaks an impasse in the understanding of this unusual group of crustaceans, providing a critical resource for testing hypotheses of adaptation to hypoxic and aphotic environments, gene family turnover, venom evolution, and the evolutionary dynamics of genome architecture. Expansion of such genomic resources is critical for deciphering the origins of numerous pancrustacean groups, with emphasis on Hexapoda.

## Materials and Methods

### Study Site

Specimens were collected from the anchialine cave system Aayin Aak (accessed via cenote Crustacea), located near Puerto Morelos, Quintana Roo, Mexico (20.7804° N, −86.9473° W). Aayin Aak is a coastal subterranean system exceeding 23 km in surveyed passage length (Devos et al. 2024) and maintains a direct hydrological connection to the Caribbean Sea. It has at least five cenotes, or surface entrances, and reaches a maximum depth of 25.6 m, with an average depth and halocline depth of approximately 15 m (Neiber et al. 2012; Anon 2025). Aayin Aak represents a stable and extreme chemosynthetic environment characteristic of remipede habitats, including permanent darkness and hypoxic and sulfide-rich waters (Neiber et al. 2012). To date, six endemic subterranean crustacean species have been reported from this cave system (Calderón-Gutiérrez et al. 2017).

### Specimen collection and preservation

Adult specimens of *Xibalbanus tulumensis* were collected by hand (Calderón-Gutiérrez) from upstream and downstream cave zone sections of Aayin Aak cave system (via cenote Crustacea) using cave diving techniques. Sampling was conducted below the halocline, within the saline (∼35 psu) water layer of the cave system. Following collection, specimens were relaxed by cooling until they ceased responding to external stimuli and promptly preserved using one of the following methods: DNA/RNA Shield (Zymo Research) or 96% ethanol. For in situ preservation, 50 ml sterile syringes with Luer 2-and 3-way stopcock valves prefilled with sterile water were triple-rinsed with cave water at the sampling site and depth prior to sampling. An individual of *X. tulumensis* was collected per syringe and immediately preserved *in-situ* by removing most of the cave water without damaging the organisms, followed by adding DNA/RNA Shield (Zymo Research) in a 9:1 ratio. Prior to adding the preservative, the sampling system was purged with DNAshield to allow a precise measurement (see ***Video S2***). All individuals were stored at −20°C until downstream processing. One specimen was deposited in Regional Collection of Invertebrates, Instituto de Ciencias del Mar y Limnología, UNAM, Mazatlán, México (ICML-EMU-14124).

### Environmental profiling

Water column profiles were measured and collected upstream and downstream from Cenote Crustacea (***Figure S4***). The upstream profile includes the surface/entrance of Cenote Crustacea to the bottom, and the downstream profile within the cave section includes the cave ceiling to the bottom. Profiles were obtained using a multiparameter sonde (YSI EXO1) equipped with temperature, conductivity, pH, and dissolved oxygen sensors. All sensors were calibrated in accordance with manufacturer specifications prior to deployment. These profiles were aligned to generate a continuous environmental profile of the system following established protocols (Calderón-Gutiérrez et al., 2023).

### DNA extraction and sequencing

For the draft assembly, high-molecular-weight DNA was extracted from the posterior body region (half of one adult organism), stored in DNA/RNA Shield with the Monarch HMW kit (New England Biolabs) following the manufacturer’s protocol, except with the proteinase K incubation time doubled. DNA quantity and purity were assessed using a NanoDrop spectrophotometer (Thermo Scientific), and fragment integrity was evaluated in a FemtoPulse fragment analyzer (Agilent, Santa Clara, CA) (***Figure S10a***). Genomic DNA was subjected to controlled fragmentation, and two HiFi libraries were prepared and sequenced independently on a PacBio Sequel II instrument at the Maryland Institute for Genome Sciences (University of Maryland; Baltimore, MD). To complement the required throughput for a genome assembly, a third run was performed using DNA extracted from the anterior region of the same individual with the Quick DNA kit (Zymo Research). Due to the fragment size distribution of the column-based purification (***Figure S10b***), the genomic DNA from this extraction was used directly for HiFi library preparation without prior shearing and was sequenced on a dedicated SMRTcell at Azenta Inc. (South Plainfield, NJ).

For chromosome-scale scaffolding, chromatin contact libraries were prepared with the Omni-C kit (Dovetail Genomics, Scotts Valley, CA) according to the manufacturer’s instructions. The whole body of individual organisms preserved in 96% ethanol were rehydrated in nuclease-free water, flash frozen with liquid nitrogen, and followed by homogenization of tissue with a pestle. Two libraries were generated from the homogenized whole body of one individual, and another four libraries were generated from the homogenized whole bodies of two pooled individuals. Cross-linking was performed by fixing the tissue with formaldehyde. The DNA was subjected to endonuclease fragmentation, end-polishing and bridge-ligation, and the resulting DNA was purified and size-selected using magnetic beads. The resulting DNA was used for Illumina library preparation via indexed adapter ligation, and the libraries were size-selected and quantified for sequencing using a TapeStation (Agilent). Sequencing was performed using a 2×150 bp paired end strategy on an Illumina NovaSeq platform at Novogene Inc.

### Genome assembly, annotation and genome size estimation

High fidelity (HiFi) long reads were assembled *de novo* using HiFiasm ver. 0.19.9-r616 (Cheng et al. 2021; Cheng et al. 2022; Cheng et al. 2024) with default parameters for haplotype purging based on read coverage. Genome heterozygosity was estimated by *k*-mer frequency profiling with KMC v3 (Kokot et al. 2017) (-k21, -ci1 -cs5000) and modeled with GenomeScope2 v2.0 (Ranallo-Benavidez et al. 2020). Chromosome-scale scaffolding was performed using Omni-C proximity ligation data. Omni-C reads were mapped to the HiFi assembly using BWA v0.7.17 (Li and Durbin 2009). The resulting BAM files were processed with the Arima Genomics Hi-C pipeline (Arima Genomics 2021) to retain inter-contig read pairs. Scaffolding process was then conducted using YaHS v. 1.2.2 (Zhou et al. 2023), and Hi-C contact maps were curated using JuicerTools implemented in Juicebox ver. 2.20 (Durand et al. 2016). Final assembly statistics, including final scaffold span and contiguity metrics were calculated using QUAST (Gurevich et al. 2013), and assembly completeness was evaluated using BUSCO (Simão et al. 2015; Manni et al. 2021) and the arthropoda_odb12 database. Repetitive elements were identified and masked prior to gene prediction.

Repeat families were first predicted *de novo* using RepeatModeler ver. Open-1.0 (Smit and Hubley 2015) with the −LTRStruct option to improve detection of long terminal retrotransposons. Then, simple, *de novo,* and database-assisted detected repeats were annotated and soft masked in a staged process with RepeatMasker Ver. 4.1.8 (Smit et al.) with the Protostomia partition of the Dfam39 database. Annotated repeats were processed with ProcessRepeats and converted to GFF3 format for downstream analysis. Repeat content was quantified by calculating the proportion of the genome masked for each family and subfamily.

Protein coding genes were predicted using Braker3 (Gabriel et al. 2024), which integrates GeneMark-ETP, AUGUSTUS and TSEBRA for evidence-based annotation, using publicly available short-read RNA-seq datasets of *Xibalbanus tulumensis* (SRA accessions SRR4113501, SRR4113500, SRR1653240, and SRR5483223). To improve annotation completeness, a broad arthropod protein database (Arthropoda.fa (Benson et al. 2015; Clark et al. 2016)) was incorporated as additional evidence with the *--prot_seq* option. Predicted proteins were functionally annotated using InterProScan v. 5.65-97.0 (Jones et al. 2014), and annotation results were merged and integrated into the BRAKER-generated GFF3 file using AGAT (*agat_sp_manage_functional_annotation.pl*) to add domain, GO term, and pathway information for downstream analyses.

### Protein annotation and evaluation

Protein-coding gene models predicted from the repeat-masked genome assembly were functionally annotated using the OmicsBox v.3.5.4 (Anon 2019) pipeline. Amino acid sequences derived from BRAKER3 gene predictions were used as input for downstream analyses. Functional inference was initially performed using BLASTp similarity searches against the NCBI non-redundant (nr) protein database implemented through OmicsBox CloudBlast (NCBI Blast+ 2.17.0 (Altschul et al. 1990)) with the following parameters: e-value = 1e-5, taxonomy filter: Arthropoda (taxid: 6656), HSP length cutoff =33, and HSP-hit coverage cutoff = 80*%*. Top-scoring hits were retained for downstream functional mapping. Taxonomic distributions of BLASTp hits were also examined to assess whether annotation patterns were broadly consistent with expected arthropod affinities (***Figure S11***). To complement homology-based annotation, conserved protein domains and functional motifs were identified using InterProScan *v.* 5.72-103.0 (Jones et al. 2014), querying against all available databases except PANTHER, which was excluded due to compatibility constraints within the cloud-based analysis environment. Gen*e* Ontology (GO) terms were assigned using the OmicsBox GO mapping and annotation pipeline by integrating BLAST results and InterProScan results using default settings. Annotated GO terms were subsequently categorized into Biological Process, Molecular Function and *C*ellular Component *(**Figure S12***).

To investigate functional categories of interest, annotated transcripts were queried based on GO identifiers and keyword searches associated with environmental sensing and signaling, ion transport and osmoregulation, oxidative stress and detoxification, reproductive processes, and venom-related genes (***Table S7***). Unique gene models were used for summary counts, while transcript-level annotations, including isoforms, were retained for visualization of scaffold-level distributions (***Figure S1***). To further improve orthology-based functional classification, protein sequences were additionally annotated using EggNOG-mapper v.2.1.13 (Huerta-Cepas et al. 2019). EggNOG annotations were used to complement BLASTp and InterProScan results and to recover additional functional assignments and KEGG Orthology (KO) identifiers (Kanehisa 2000).

KEGG pathway annotation was performed by integrating KO assignments derived from EggNOG mapper and OmicsBox. KO terms were mapped to reference metabolic and signaling pathways using the KEGG database (Kanehisa 2000) to reconstruct pathway-level functional profiles. Annotated pathways (***Table S8***) were summarized across major functional categories to assess broad metabolic capacity (***Figure S3***). Targeted pathways of interest, including hypoxia-related signaling (e.g., HIF-1 signaling pathway) and core metabolic processes such as glycolysis/gluconeogenesis, were examined in detail by mapping identified KO terms onto KEGG reference diagrams to evaluate pathway completeness and gene representation (***Figures S5–S6***).

### Maximum Likelihood Phylogenetic Reconstruction and Divergence Time Estimation

To infer a dated phylogenomic tree for comparative analyses, BUSCO genes were extracted from the peptide files of 22 arthropod genome projects (20 pancrustaceans, one myriapod, and one chelicerate), including *X. tulumensis* (BUSCO v.5.8.2 with the arthropoda_odb10 dataset (Manni et al. 2021)). BUSCO genes were binned as fasta sequences using a custom Python script and multiple sequence alignments were performed using mafft v.7 (Katoh 2002; Katoh and Standley 2013) with default parameters, followed by trimming with Gblocks v0.91b (Castresana 2000) (- b4=10 −b5=n). A total of 225 alignments with at least 16 taxa (>70% complete) were retained and concatenated using phyutility (Smith and Dunn 2008).

Divergence times were estimated with MCMCtree v4.9e (dos Reis and Yang 2019) (PAML software package (Yang 2007)) based on the 225 alignments and supplying a constraint phylogeny based on dense sampling of Pancrustacea (Bernot et al. 2023). Six temporal constraints derived from established benchmarks were incorporated (***Table S9***). To facilitate computational efficiency, the approximate likelihood method was employed, involving the calculation of a Hessian matrix in codeml (Yang 2007) using empirical base frequencies and the LG+G5 substitution model (LG with five Gamma-distributed rate categories). Posterior distributions were estimated using a correlated molecular clock model. Four independent MCMC chains were executed for 12 million generations each, with the initial 40,000 generations discarded as burn-in. Convergence was assessed by plotting the node ages of independent runs against each other in R (R Core Team 2025).

Phylogenomic dating recovered an estimated median divergence time between *X. tulumensis* and Hexapoda of 477 Ma, whereas previous hypotheses have suggested older dates of approximately 519 Ma (Schwentner et al. 2017) or exceeding 540 Ma in one study (Bernot et al. 2023). These discrepancies in median node ages are attributable to differences in taxonomic sampling as well as the use of node calibration priors (specifically, the root age prior, which we set conservatively in this study). The resulting dated tree was used to provide a temporal context for downstream analyses of gene family evolution.

### Gene Family Evolution

To identify orthologous groups, proteomes from 22 taxa (including *X. tulumensis*) were analyzed using OrthoFinder v2.5.4 (Emms and Kelly 2019). To eliminate redundant transcript isoforms, CD-HIT v.4.6 (Li and Godzik, 2006) was executed on each dataset with a 98% sequence identity threshold, retaining only the single longest isoform per locus. To minimize noise from lineage-specific repetitive elements, genes annotated as Transposable Elements (TEs) were removed from the dataset, and the orthology inference was re-calculated on the filtered data. Functional annotation was performed by selecting the longest sequence per orthogroup as a representative for analysis with eggNOG-mapper v2.1.12. Gene family expansions and contractions were modeled using CAFE5 v1.1 (Mendes et al. 2021). Following "best practices" outlined in the CAFE5 documentation, stringent filtering was applied to the Hierarchical Orthologous Groups (HOGs); specifically, HOGs with a variance of over 50 genes between the most and least speciose taxa were excluded, as were lineage-specific HOGs present in only one species. Several models of gene family evolution were evaluated, including Uniform and Poisson distributions for the root state, as well as two gamma rate categories paired with Poisson distributions. To account for potential assembly or annotation artifacts, an error model was estimated under two configurations: root Uniform and root Poisson. Significant expansions and contractions within the *X. tulumensis* lineage were identified using a significance threshold of p < 0.05.

Targeted searches of genes were performed using BLASTp, with sequences from the *Drosophila melanogaster* and *Daphnia pulex* proteomes used as queries for Hox and opsin gene families, respectively. For Hox gene surveys, BLASTp searches were performed for *X. tulumensis*, seven other crustacean species with high-quality genomes, and representative myriapod and tick as outgroups (***Figure 2c***). The top hits with e-value < 1 x 10e-20 were retained and multiple sequence alignment was performed using CLUSTAL-Omega, followed by inference of the gene tree using IQ-TREE v.2 (Nguyen et al. 2015; Minh et al. 2020) with an LG+I+G substitution model and 1000 ultrafast bootstrap replicates (***Figure S13***). For the opsin analysis, the single opsin sequence identified in the *X. tulumensis* genome was embedded in a 631-terminal multiple sequence alignment of crustacean opsins (Palecanda et al. 2022). Alignments and gene trees are provided as supplementary materials.

Identification of miRNA families was performed using MirMachine (Umu et al. 2023) for the same nine taxa included in the Hox gene analysis (***Table S3***). A combined model (protostome and deuterostome) search was implemented and Pancrustacea was specified as the search node. The distribution of filtered, high-quality hits was mapped across taxa, after ordering miRNA families by phylostratigraphic depth (***Figure 2c***).

### Evolutionary Analysis of Venom-Related Gene Families

To explore the evolutionary dynamics of venom components, CAFE5 (Mendes et al. 2021) results were intersected with a manually curated database of putative toxins. Reference sequences for toxin families were retrieved from von Reumont et al. (2017) and subjected to a Blastp search against the *X. tulumensis* proteome. High-confidence orthology was ensured by applying stringent filtering criteria, retaining only hits with 100% identity and an E-value < 1 x 10^-80^. These identifiers were subsequently mapped to their respective HOGs to identify venom families undergoing significant expansion or contraction. For HOGs identified as significantly expanded in

*X. tulumensis*, signatures of episodic positive selection among paralogs were tested using the adaptive Branch-Site Random Effects Likelihood (aBSREL) method in HyPhy v2.5 (Kosakovsky Pond et al. 2020). Nucleotide sequences were retrieved from the genome assembly and codon-aligned using PAL2NAL v14 (Suyama et al. 2006), guided by the corresponding multiple sequence alignments of the amino acids. Gene trees were reconstructed using the amino acid sequences and IQ-TREE (Nguyen et al. 2015) implementing ModelFinderPlus (Kalyaanamoorthy et al. 2017) and Ultrafast Bootstrap Replicates (Hoang et al. 2018). Finally, "orphan" HOGs (those unique to *X. tulumensis*) that contained matches to known toxin families were identified to highlight potential lineage-specific venom innovations.

### Macrosynteny

For comparative analyses of macrosynteny, genomic positions of orthologs belonging to Bilaterian-Cnidarian-Sponge (BCnS) ancestral linkage groups (ALGs) defined by Simakov et al. (2022) were identified in the assembly of *Xibalbanus tulumensis* and visualized in Oxford dot plots using odp v.0.3.0 (Schultz et al. 2023). Orthologs were inferred between species by finding reciprocal-best protein matches using *Diamond* v.2.2 (Buchfink et al. 2015) and OrthoFinder v.2.3.7 (Emms and Kelly 2019). The reciprocal-best hits were used to identify macrosyntenic chromosomes between species by performing Bonferroni-corrected one-sided Fisher’s exact tests (Simakov et al. 2020). We used in-built tools in odp v.0.3.0 to visualize syntenic patterns as Oxford dot plots and ribbon plots, excluding scaffolds less than 500 Mb.

To infer the evolutionary history of linkage groups across the phylogeny, the same ALG set was identified in the genomes of four insects (*Entomobrya proxima* [GCA_029691765.1]; *Tomocerus qinae* [GCA_020055645.1]; *Thermobia domestica* [GCA_964235325.1]; *Ischnura elegans* (GCA_921293095.2]), three Branchiopoda (*Artemia franciscana* [GCA_032884065.1]; *Lepidurus arcticus* [ENA PRJEB111934], *Daphnia pulex* [GCF_021134715.1]); one Copepoda (*Tigriopus californicus* [GCA_007210705.1]); two Communostraca (*Penaeus (Litopenaeus) vannamei* [GCA_042767895.1], *Pollicipes pollicipes* [GCA_011947565.2]); and two non-pancrustacean outgroups (the myriapod *Strigamia acuminata* [GCA_949358305.1] and the chelicerate *Ixodes scapularis* [GCF_016920785.2]). Due to the lack of a chromosome-level genome for an exemplar of Oligostraca at the time of analysis (e.g., Ostracoda; Mystacocarida; Branchiura), the deepest split within Pancrustacea could not be represented.

Fusion-with-mixing events were contextually defined as events involving a minimum of 40 genes from two linkage groups and whose distribution along a chromosome required a minimum of five swap events to explain. Mixing events were mapped for all taxa individually, followed by ancestral state reconstruction of major events on the LG + C60 + F + G maximum likelihood tree topology of Bernot et al. (2023).

## Supporting information

Xibalbanus 2026: Supplementary Figures

Xibalbanus 2026: Supplementary Tables

## Acknowledgments

Funding for this work was provided by the National Science Foundation (NSF) OCE-2136322 (E.B.), with lab work also supported by internal funds from Texas A&M University−San Antonio (A&M-SA; E.B.), Instituto de Ciencias del Mar y Limnología - Universidad Nacional Autónoma de México (ICMyL−UNAM; R.L.H.), the University of Wisconsin−Madison (UWM; P.P.S.) and Stephen F. Austin State University (SFASU; F.C.G.). Collection activities were conducted under a SEMARNAT permit to Maite Mascaró (UNAM-Sisal), associated with CONAHCyT Ciencia Básica y de Frontera project CBF2023-2024-1658 (PI: Maite Mascaró), “Efectos del cambio climático y la contaminación en las especies del acuífero de la Península de Yucatán: un enfoque ecofisiológico,” which also provided financial support for the postdoctoral work of Efraín Chávez Solís. Genomic sequencing consultation was kindly provided by Luke Tallon (Maryland Genomics), Jorge Franco and Omar Sayed (Novogene Corporation, Inc). Computational analyses were conducted using local laboratory workstations at Texas A&M University−San Antonio and computational infrastructure hosted at the Chihuil server at UNAM−Mazatlán (R.L.H.), UWM (P.P.S), the Computational Biology Core at the University of Connecticut (granted to C.E.S.L.), the Supercomputer at Western Connecticut State University (C.E.S.L.) and Kean University (J.A.B). We thank Vlada Dekina for providing the underwater photograph of the anchialine cave habitat of *Xibalbanus tulumensis.* We thank Joan Bernhard for supporting diving operations (NSF OCE-2136377), as well as Efraín Chávez Solís, Shari Rohret, Vlada Dekina, Natalie Gibb and Under the Jungle for field support, and the landowner for providing site access to Cenote Crustacea (Aayin Aak system) in Puerto Morelos.

## Author contributions

R.L.H., P.P.S., E.B., F.C.G., and B.Y.R. conceptualized the study. E.B., P.P.S., C.S.L., R.L.H., and F.C.G. wrote the manuscript. E.B. and P.P.S. contributed equally to this work and share first authorship. E.B., R.L.H., P.P.S., and F.C.G. secured funding for the study. F.C.G. collected specimens and environmental profiles. N.S. provided fieldwork logistics. R.L.H. and P.P.S. extracted DNA and prepared sequence libraries. A.G.B. and R.L.H. assembled the genome and performed repeat and gene annotations, with assistance from C.S.L. and E.B. P.P.S. performed Hox, miRNA, and opsin analyses. C.S.L. performed HOG and aBSREL analyses. J.A.B. and P.P.S. performed macrosynteny analyses. P.P.S. prepared the graphical abstract. All authors discussed the results, contributed to, and reviewed the final manuscript.

## Data and code availability

Raw sequencing data generated in this study have been deposited in the NCBI Sequence Read Archive (SRA) under BioProject accession PRJNA1456311. Associated BioSample accessions include SAMN57478728, SAMN57478729 and SAMN57404124, and raw sequence datasets are available under SRA accessions SRR38289187−SRR38289189 and SRR38287356−SRR38287361. The chromosome-level genome assembly has been deposited in GenBank under accession XXXX00000000 (*Under Review*).

Supplementary videos and datasets, annotations, and additional project files are available through Zenodo: 10.5281/zenodo.18918102.

Environmental data associated with the Aayin Aak System sampling site are available through BCO-DMO: Project: 868765.

Additional supplementary figures and images are available through FigShare: a) BCnS synteny analyses, 10.6084/m9.figshare.34863156; and b) Image silhouettes (***Figure 4b***), 10.6084/m9.figshare.34872654.

