## Supplementary material for "Genomic expansion in the dark: A repeat-rich genome of a cave-dwelling remipede sheds light on the evolution of the hexapod sister group": Xibalbanus 2026: Supplementary Figures

**Video S1**: ***Xibalbanus tulumensis*** **swimming and prey capture behavior.** (See Zenodo: Video S1)

**Video S2**: ***In situ*** **DNA/RNA shield preservation of *Xibalbanus tulumensis*.** (See Zenodo: Video S2)

**Figure S1. Chromosome-level scaffold distribution of select functional gene categories.**
Heatmap of Z-score normalized transcript counts across select functional categories: **a)** osmoregulation (osmo; 205 unique gene models), **b)** oxidative stress and detoxification (stress; 62), **c)** environmental sensing and signaling (environ; 88), **d)** reproductive processes (reprod; 12), and **e)** toxin and venom associated enzymatic activity (venom; 521), per chromosome-level scaffold (***Table S7***). Heatmap values are based on transcript-level annotations and therefore include multiple isoforms from the same gene model where present. Cooler colors indicate relative relatively higher transcript counts and warmer colors indicate relative depletion within each functional category among scaffolds.

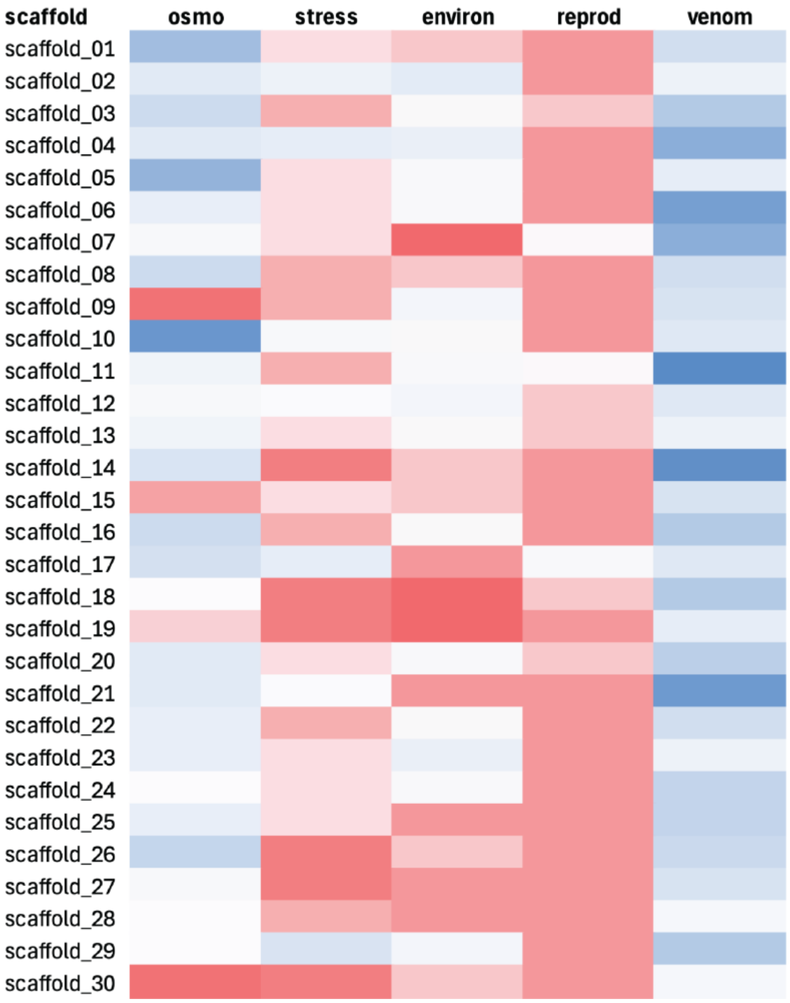

**Figure S2. Distribution of InterProScan superfamilies among proteins lacking BLAST similarity for *Xibalbanus tulumensis*.**

All sequences (n=1,418) were assigned to conserved domain families based on InterProScan and associated GO annotations despite lacking sequence-level homology. The "others" category represents a collection of lower-frequency domain families, indicating broad functional diversity among highly diverged proteins.

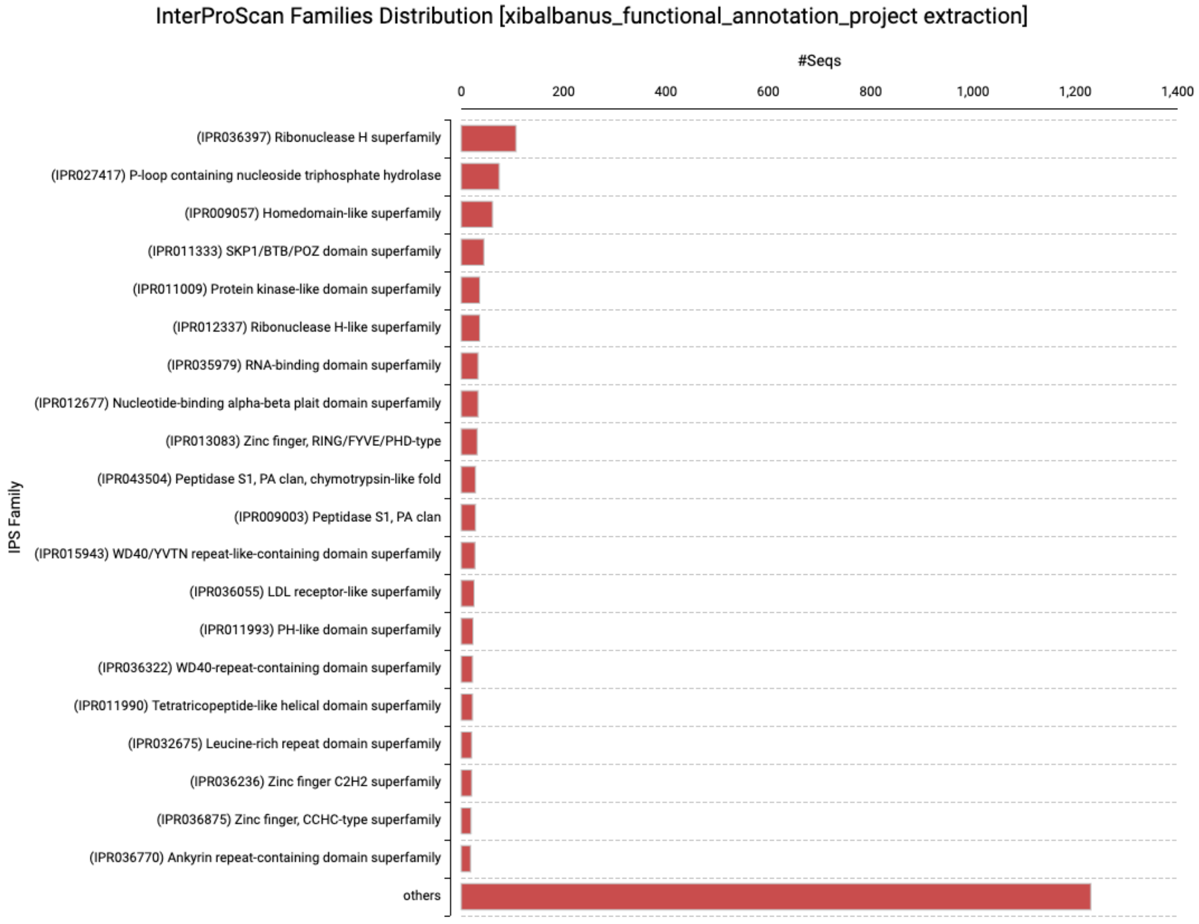

**Figure S3. Distribution of KEGG pathways across major functional categories in *Xibalbanus tulumensis*.**

KEGG annotation identified 310 pathways spanning five major functional categories, including Metabolism (n=116 pathways), Organismal Systems (n=92 pathways), Environmental Information Processing (n=39 pathways), Cellular Processes (n=33 pathways), and Genetic Information Processing (n=30 pathways). No single category exhibited disproportionate enrichment.

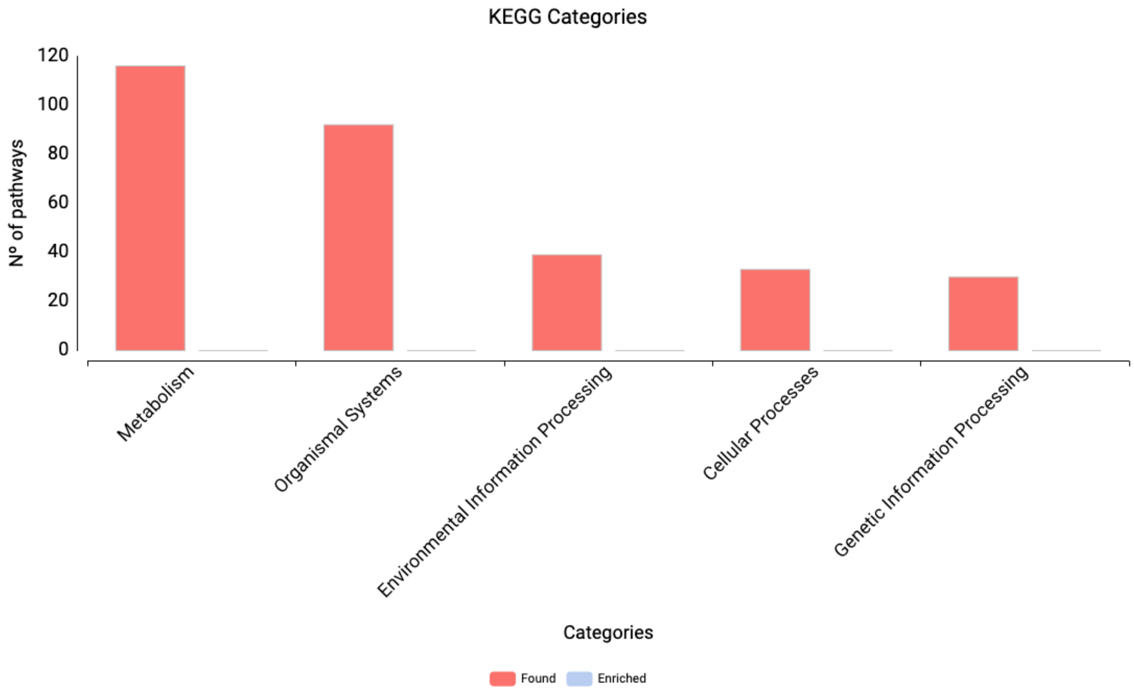

**Figure S4. Environmental profiles of Aayin Aak system via cenote Crustacea (Puerto Morelos, Quintana Roo, Mexico).**

Vertical profiles of salinity (blue), temperature (red), pH (brown), and dissolved oxygen (DO; green) across depth (m) for **a)** downstream and **b)** upstream from cenote/cave entrance. Profiles show pronounced physicochemical stratification, including two haloclines with increasing salinity at depth and corresponding shifts in temperature and pH. Dissolved oxygen varies strongly with depth, with reduced concentrations (<5%) downstream, where *Xibalbanus tulumensis* is most abundant, and comparatively higher, though variable, oxygen availability upstream in the saline layer (~35 psu).

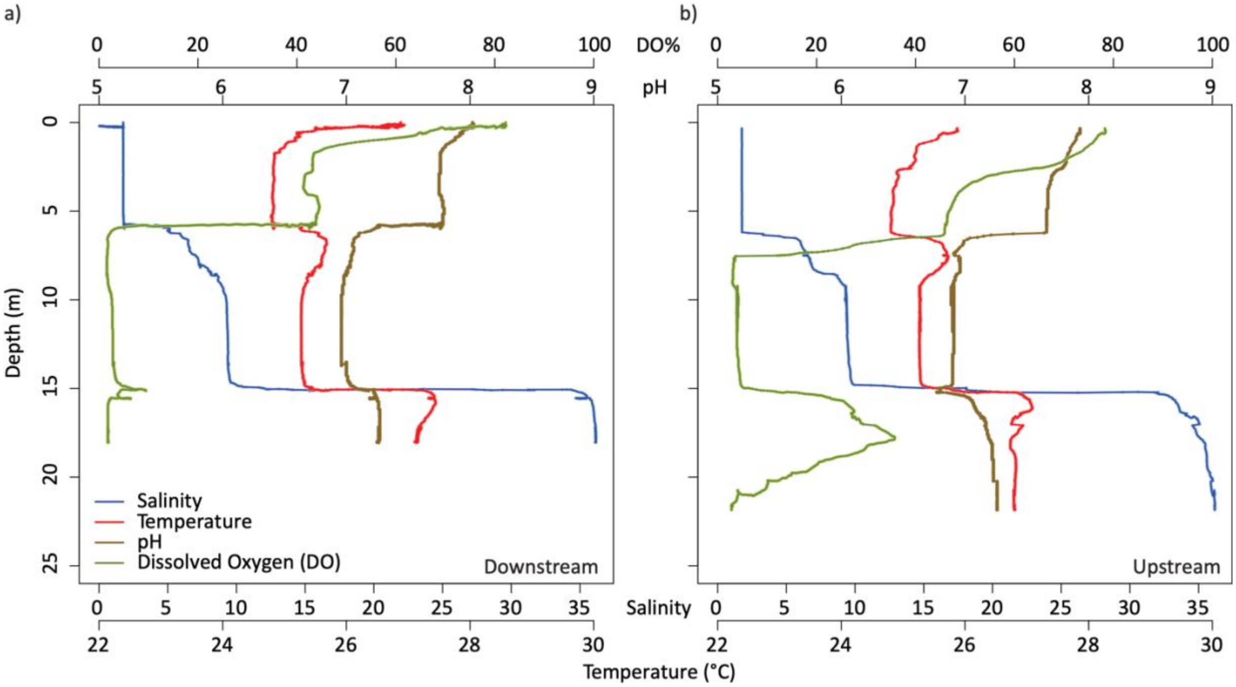

**Figure S5. Hypoxia-inducible factor (HIF-1) signaling pathway in *Xibalbanus tulumensis*.**

KEGG pathway map (ko04066) showing components of the HIF-1 signaling pathway identified in the *X. tulumensis* genome. Colored boxes indicate detected orthologs, including regulators of oxygen sensing (EGLN/PHD, VHL complex), transcriptional activation (HIF1A, EP300), and upstream signaling pathways (e.g., PI3K–AKT, MAPK, mTOR). Downstream targets involved in metabolic reprogramming are also present, including key glycolytic enzymes (e.g., HK, PFK, ALDO, PGK, LDH). These results support conservation of the canonical hypoxia response pathway without evidence of broad gene family expansion.

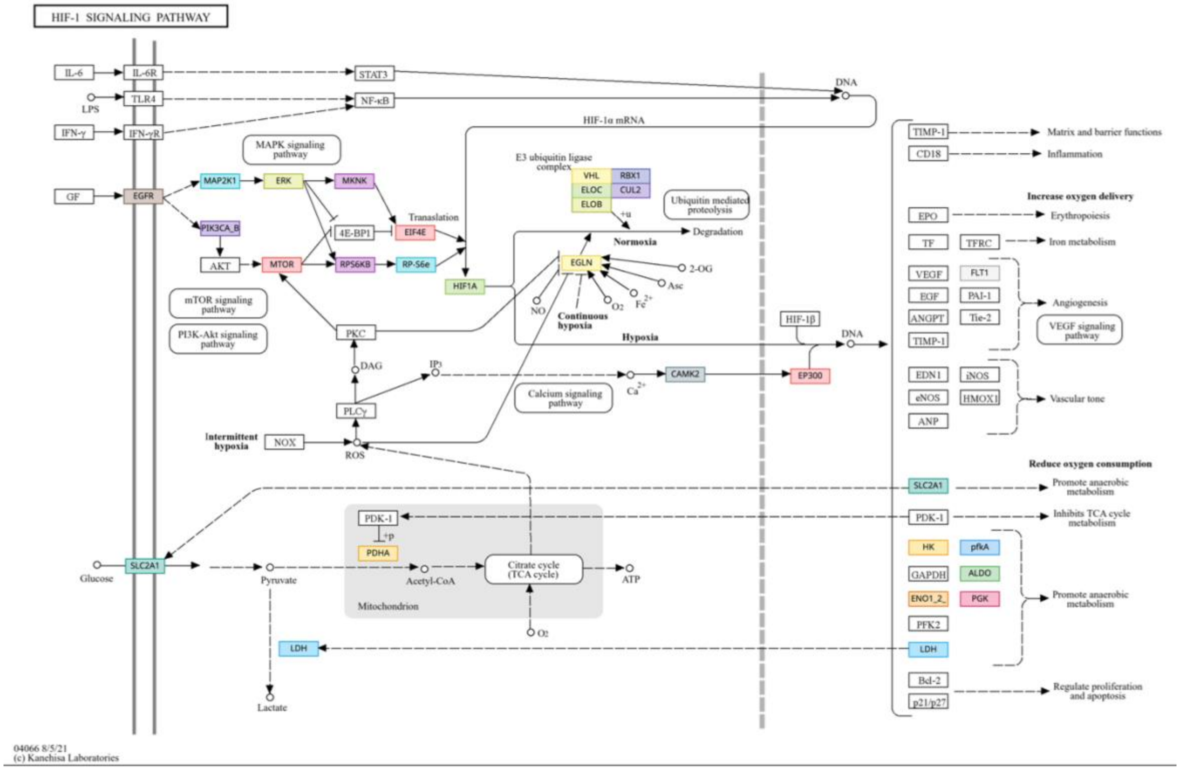

**Figure S6. Glycolysis and gluconeogenesis pathway in *Xibalbanus tulumensis*.**

KEGG pathway (ko00010) map showing the distribution of annotated enzymes in *X. tulumensis*. Colored boxes indicate enzymes identified in the genome, while gray boxes represent pathway components not detected. Core glycolytic enzymes are broadly represented, including key regulatory steps (e.g., hexokinase, phosphofructokinase, and pyruvate kinase) and downstream components involved in anaerobic metabolism (e.g., lactate dehydrogenase).

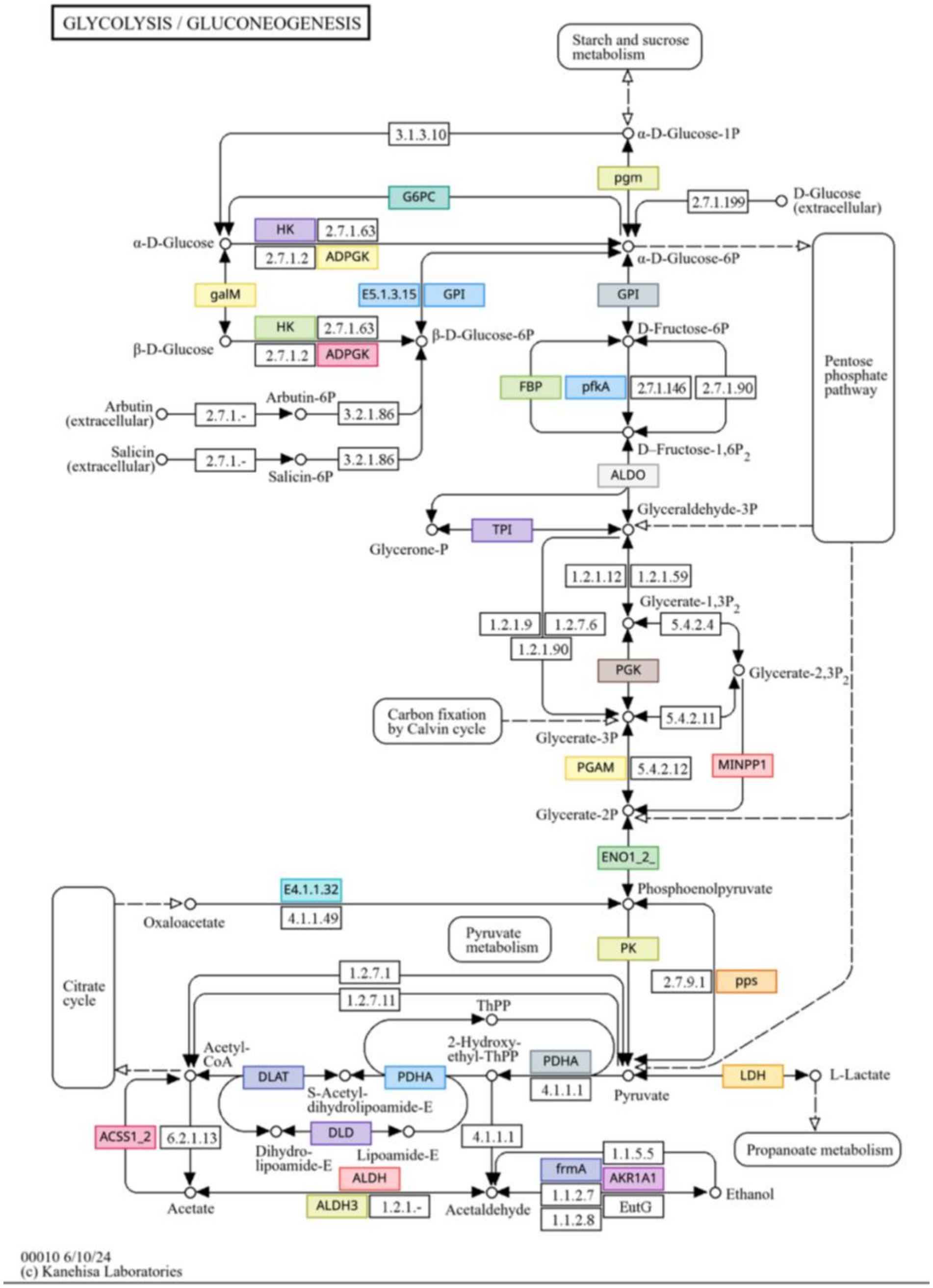

**Figure S7. Gene family expansion and contraction across pancrustacean taxa under alternative CAFE models.**

Gene family expansions (green) and contractions (red) inferred across 22 pancrustacean taxa under seven alternative CAFE models, including Poisson, PoissonK2, Uniform, PoissonK1, UniformE, PoissonK1E, and PoissonE. Bar lengths represent the number of expanding and contracting hierarchical orthologous groups (HOGs) along each lineage. Overall patterns of gene family turnover are broadly consistent across models, with similar distributions of expansion and contraction observed among taxa.

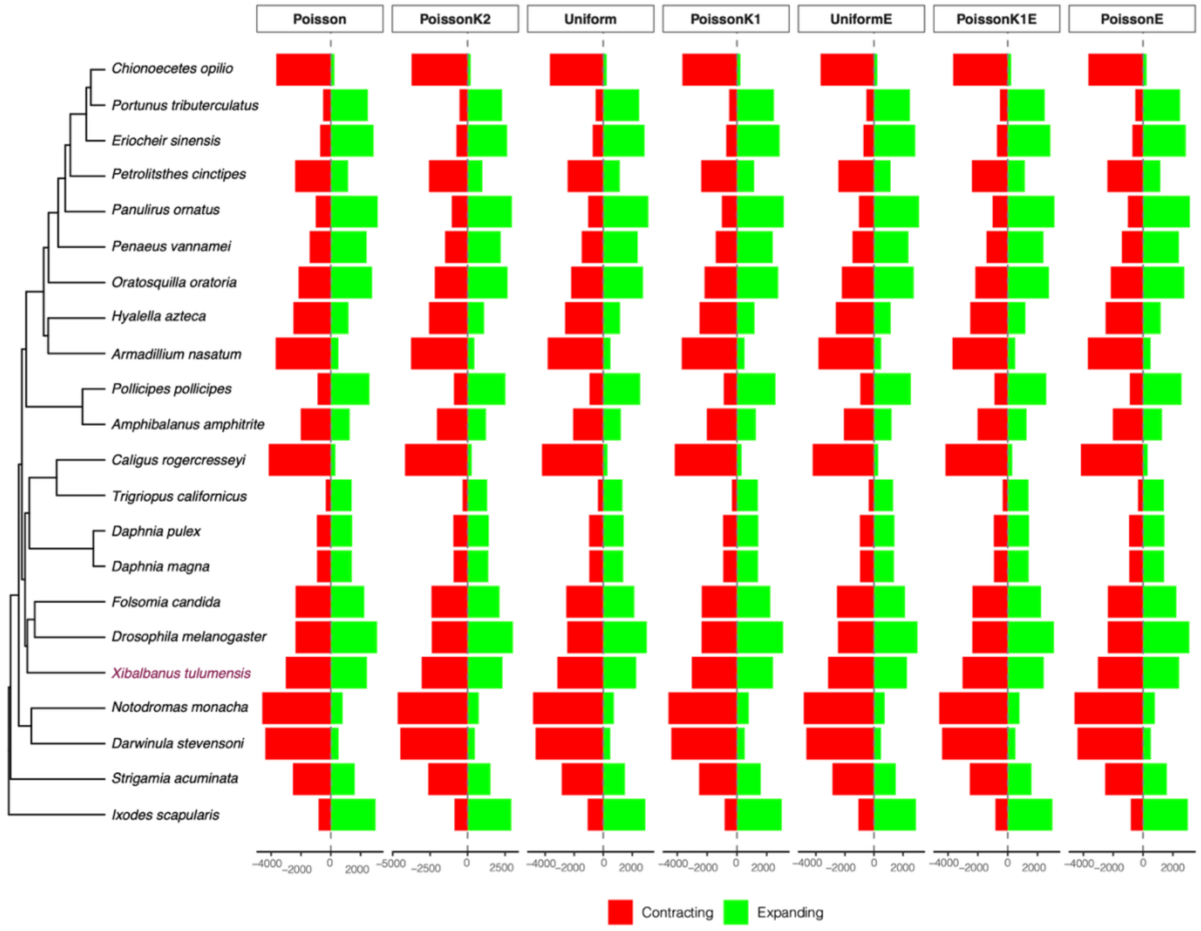

**Figure S8. Macrosynteny between *Xibalbanus tulumensis* and an expanded sampling of representative pancrustacean genomes.**

Chromosome-scale scaffolds are shown for thirteen species, with ribbons connecting homologous genomic regions identified by conserved orthologs. Pancrustacean taxa include *Pollicipes pollicipes, Penaeus (Litopenaeus) vannamei, Artemia franciscana, Lepidurus arcticus, Daphnia pulex, Tigriopus californicus, Xibalbanus tulumensis, Entomobrya proxima, Tomocerus quinae*, *Thermobia domestica,* and *Ischnura elegans*; *Ixodes scapularis* and *Strigamia acuminata* were included as arthropod outgroups.

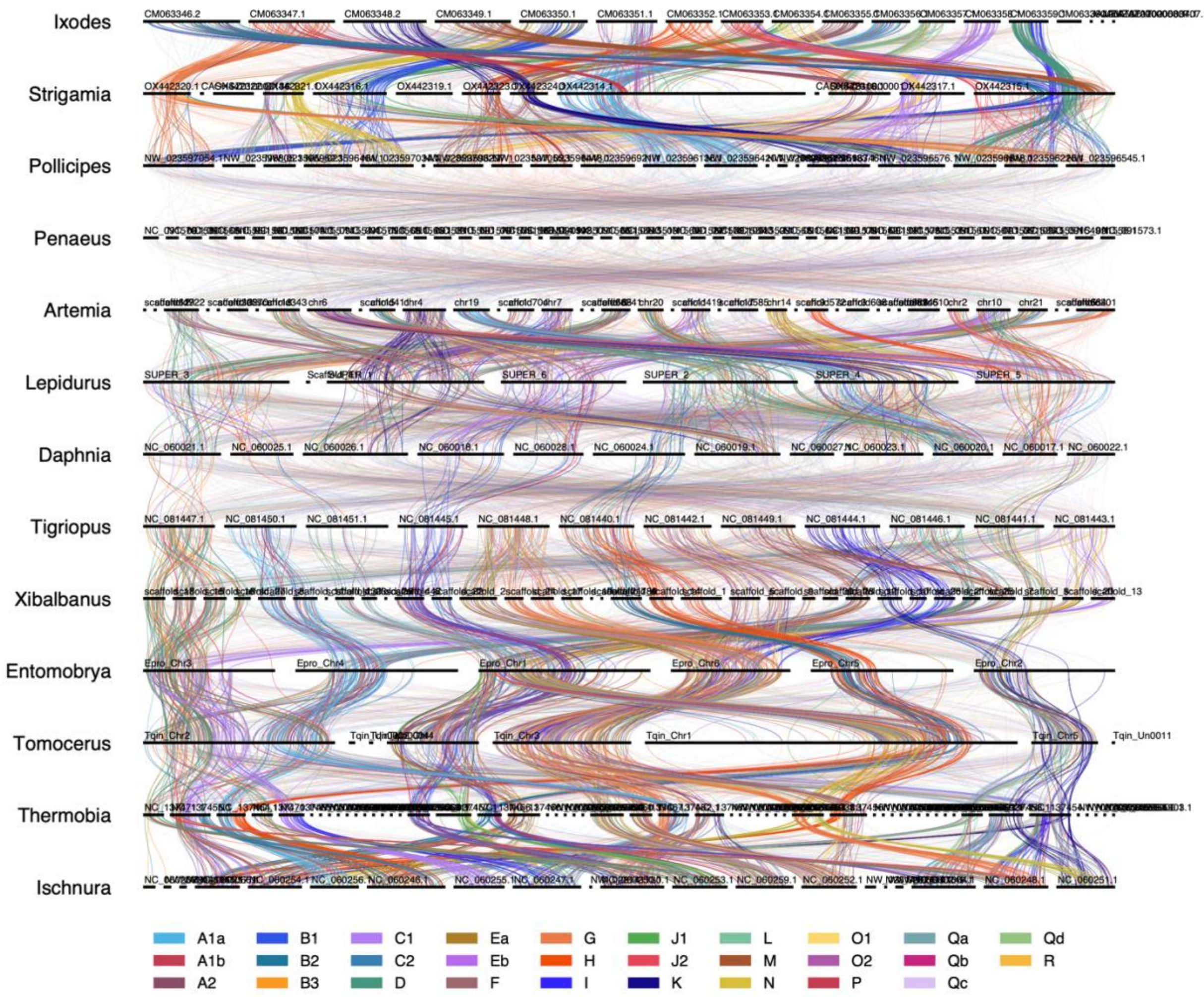

**Figure S9. Oxford dot plots showing correspondence between BCnS ancestral linkage groups and select crustacean genomes.**

**a)** *Penaeus (Litopenaeus) vannamei* and **b)** *Daphnia pulex*. Reciprocal best DIAMOND BLASTP matches were used to identify orthologous gene pairs between each genome and the 29 Bilaterian–Cnidarian–Sponge ancestral linkage groups (BCnS ALGs; Simakov et al. 2022).

**a)** ***Penaeus (Litopenaeus) vannamei***

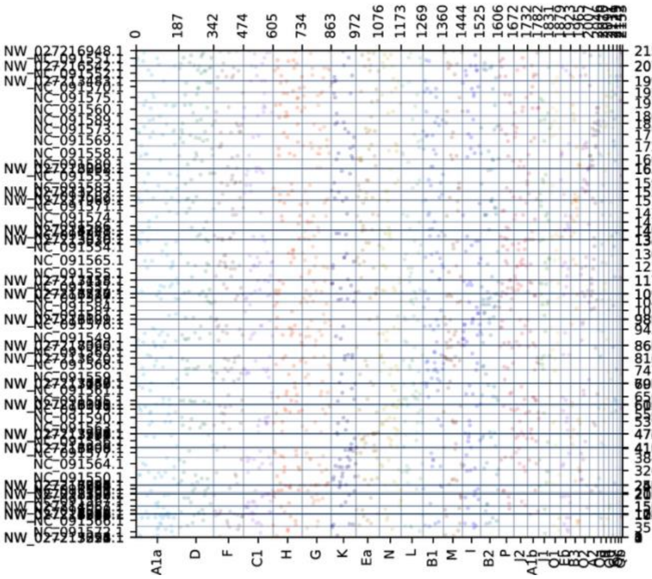

**b)** ***Daphnia pulex***

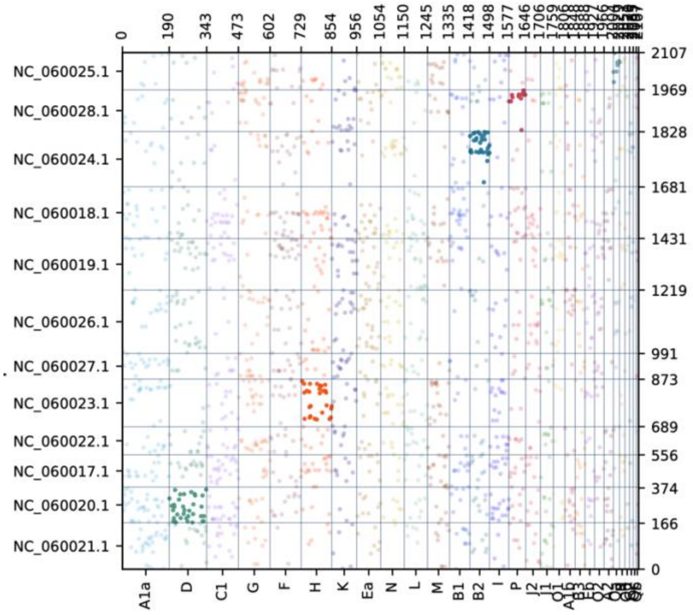

**Figure S10. Fragment-size distributions of genomic DNA of *Xibalbanus tulumensis*.**

**a)** High molecular weight (HMW) DNA from the posterior region and **b)** column-purified genomic DNA from the anterior region.

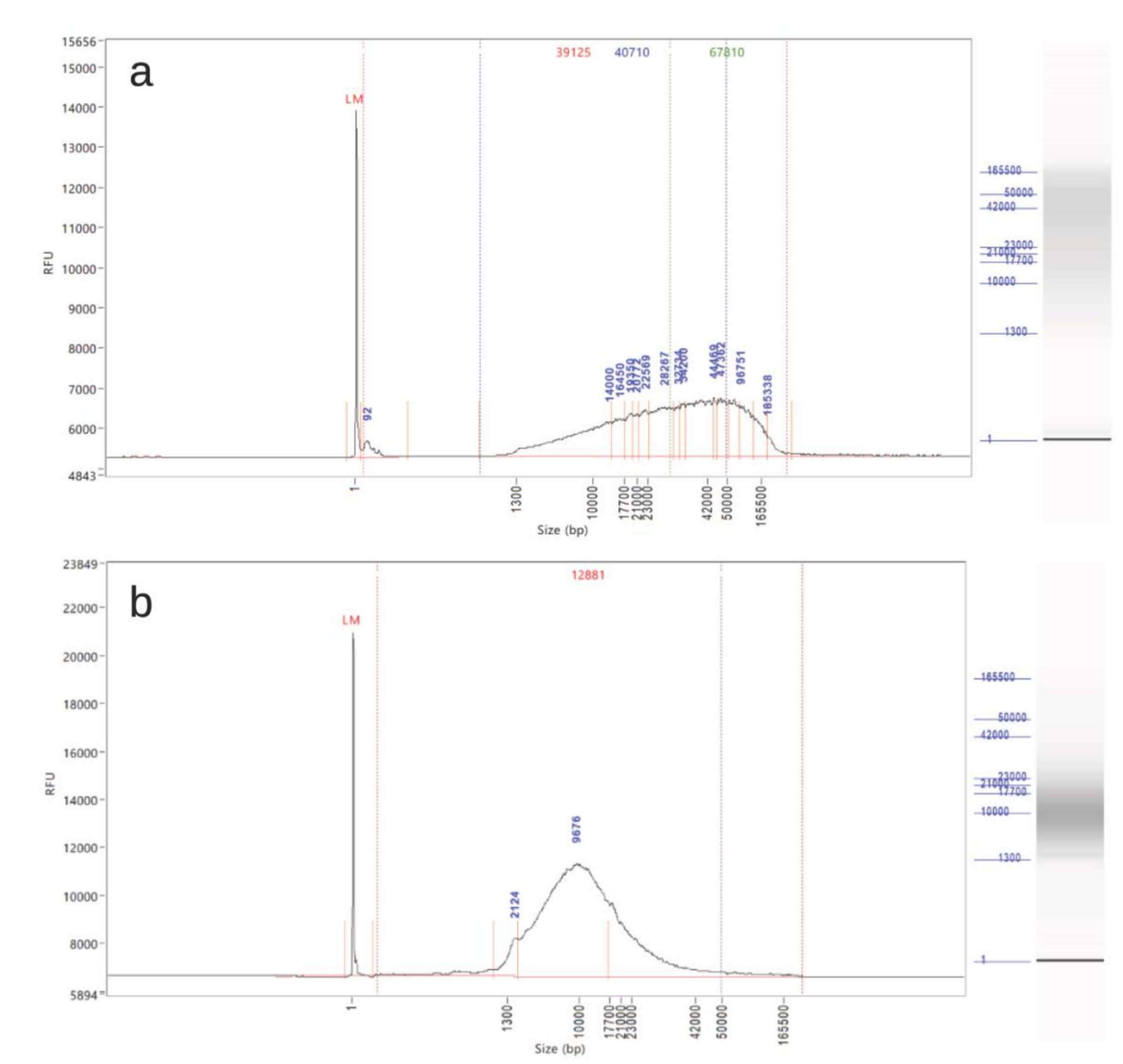

**Figure S11. Taxonomic assignment of predicted proteins based on BLASTp similarity for *Xibalbanus tulumensis*.**

Top hits were dominated by insect taxa, particularly Blattodea and Orthoptera, with additional matches to chelicerates and diverse crustacean lineages.

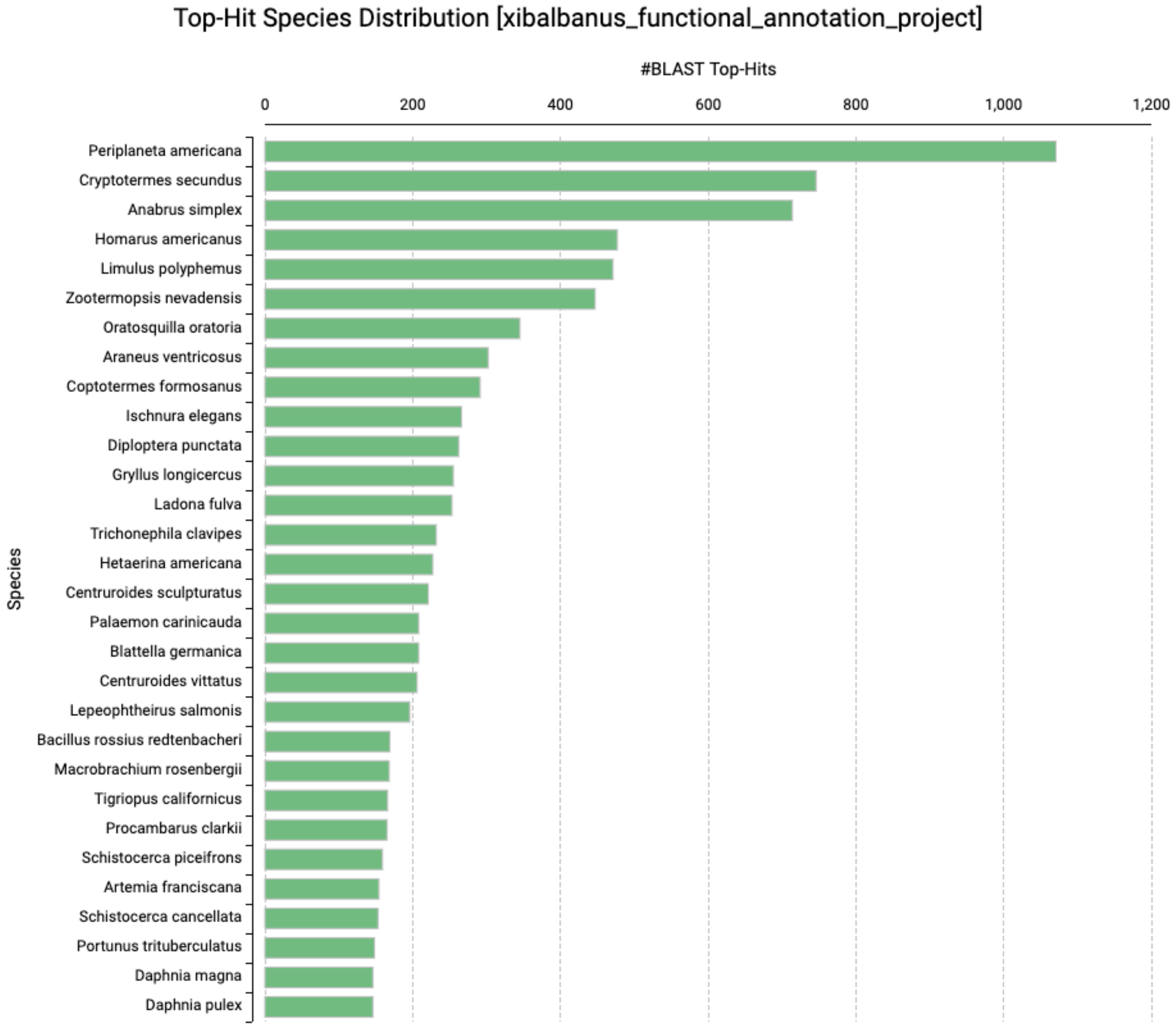

**Figure S12. Gene Ontology (GO) classification of predicted proteins in *Xibalbanus tulumensis.***

GO annotations were assigned to predicted protein-coding genes using sequence similarity and domain-based approaches in OmicsBox. Bar plots show the distribution of annotated genes across Level 2 GO categories for the three main ontologies: **a)** Biological Process, **b)** Molecular Function, and **c)** Cellular Component. The number of genes assigned is indicated on the y-axis.

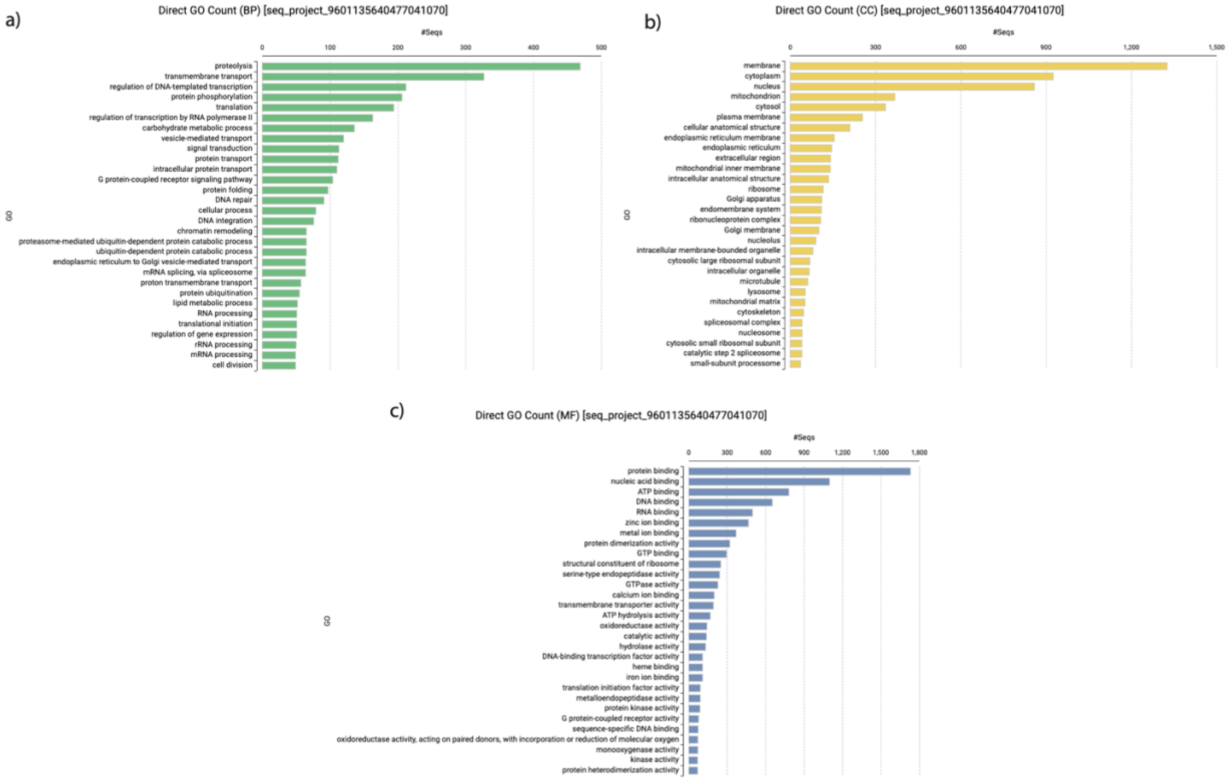

**Figure S13. Phylogenomic organization of Hox gene clusters across pancrustacean taxa.** Phylogenetic relationships are shown on the left (IQ-TREE). Colored arrows indicate the relative order and transcriptional orientation of Hox genes. While most taxa retain a conserved, colinear Hox cluster, *Xibalbanus tulumensis* exhibited a fragmented arrangement across scaffold 17, with several Hox genes not recovered in the current genome assembly.

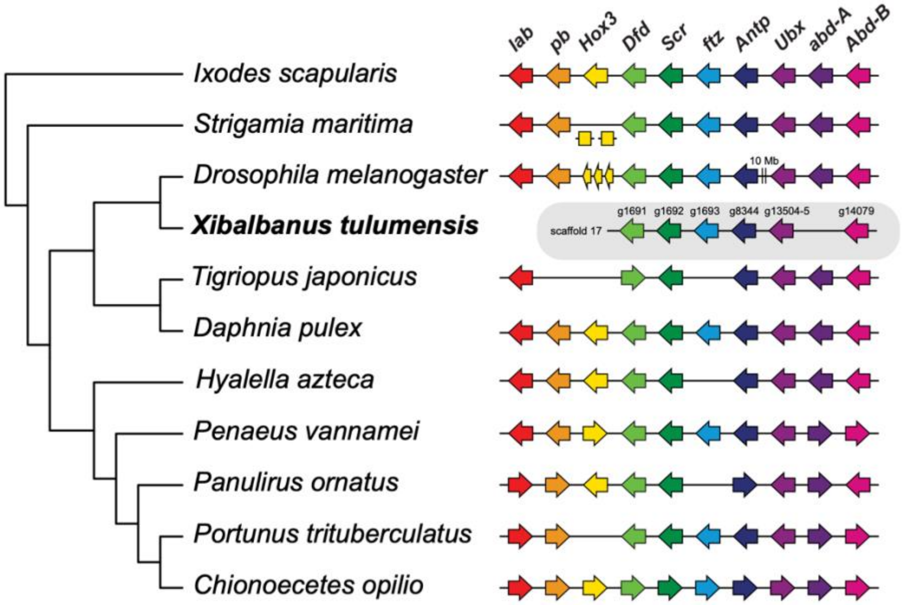
